# Lineage 2–Beijing *Mycobacterium tuberculosis* strains suppress BCG-trained innate immunity early after infection

**DOI:** 10.64898/2026.03.29.714649

**Authors:** Naomi J Daniels, Todia Setiabudiawan, Taru S Dutt, Philip C Hill, Marcela Henao-Tamayo, Reinout van Crevel, Joanna Kirman

## Abstract

Genetically distinct lineages of *Mycobacterium tuberculosis* differ in their virulence, transmissibility, and immune evasion capacity. The modern Lineage 2-Beijing (L2-B) clade of *Mtb*, which is highly prevalent in Asia and now globally distributed, is associated with resistance to the bacille Calmette Guérin (BCG) vaccine. Using a murine BCG vaccination model followed by aerosol challenge with geographically matched L2-B or Lineage 4 clinical isolates, we examined how L2-B circumvents vaccine-induced immunity. BCG-trained alveolar macrophages showed impaired control of L2-B growth *in vitro*. Spatial transcriptomic profiling of infected lungs revealed broad transcriptional reprogramming, including selective suppression of pathways essential for macrophage activation and trained innate immunity. Similar immune dysregulation was observed in peripheral blood from BCG-vaccinated household contacts of L2-B smear-positive TB patients. Together, these findings suggest mechanisms of lineage-specific immune evasion in TB, highlight key components of BCG-induced protection, and support re-evaluation of current TB vaccine strategies.

## Introduction

Tuberculosis (TB), caused by *Mycobacterium tuberculosis* (*Mtb*), is a leading cause of infectious disease deaths globally, with an estimated 1.23 million deaths in 2024^1^. Genetically distinct lineages of *Mtb* differ in virulence, immunogenicity, and vaccine responsiveness^2,3^. The global spread of the modern Lineage 2-Beijing (L2-B) clade of *Mtb* has been associated with increased transmissibility, drug resistance, higher relapse rates, and greater capacity for immune evasion than non-L2-B strains^4,5^.

Epidemiological and experimental evidence suggest that L2-B strains can evade immunity induced by the TB vaccine, bacille Calmette-Guérin (BCG). In a case-contact study in Indonesia, BCG vaccination protected against infection with non-L2-B *Mtb* strains but failed to protect against infection with L2-B strains, with adjusted RR for interferon gamma release assay (IGRA) conversion 0.4 (95% CI 0.27–0.61) and 1.02 (95% CI 0.56–1.85), respectively^6^. In murine models, BCG-vaccinated mice had reduced protection against the L2-B strain HN878^7,8^ and clinical L2-B isolates^9^, compared to protection against non-L2-B strains.

It remains unclear which components of BCG-induced responses are essential for protection, and which host pathways are targeted during evasion by L2-B strains. While much research has focused on adaptive immunity induced by BCG, less is known about the effect of BCG on the innate response that may be critical for early control of *Mtb* infection. BCG is a potent inducer of trained immunity: long-lasting functional reprogramming of innate immune cells that enhances antimicrobial responses^10,11^, driven by epigenetic and metabolic changes in monocytes and macrophages, including chromatin remodelling, glycolytic reprogramming, and enhanced cytokine production^12–15^. The effects of innate immune training are likely to be most critical at the primary site of *Mtb* infection: the lung.

Alveolar macrophages (AM) are critical for the early immune containment of inhaled *Mtb*^16^. These long-lived tissue-resident cells can either restrict or permit *Mtb* growth depending on their activation state and metabolic profile^17,18^. Therefore, the AM state may determine whether the host achieves early control or progresses to active disease^19^. BCG vaccination reshapes the lung macrophage compartment, expanding a population of CD11b^hi^ alveolar macrophages^20^ that may be more functionally activated, potentially enhancing early responses to infection. Whether L2-B strains can evade trained immunity induced in the lung following BCG vaccination is unknown.

In this study, we investigate how L2-B evades BCG-mediated immunity using a murine model of aerosol *Mtb* challenge. Using geographically matched clinical *Mtb* strains: L2-B and Lineage 4 (L4), we focused on the early stages of *Mtb* infection in the BCG-vaccinated mouse lung to identify transcriptional and metabolic programs that were selectively suppressed by L2-B. Similar immune pathways dysregulated in the infected mouse lung were discovered in the peripheral blood of BCG-vaccinated human household contacts of L2-B smear-positive TB patients, revealing host pathways targeted by L2-B *Mtb*. Together, these findings identify a conserved, lineage-specific immune evasion strategy whereby L2-B *Mtb* escapes immune control and point to facets of BCG-induced immunity that are vital for protection and vulnerable to immune evasion.

## Results

### CD11b^hi^ alveolar macrophages dominate early *Mtb* infection in BCG-vaccinated lungs and mediate strain-dependent growth restriction

Mucosal BCG immunisation expands CD11b^hi^ alveolar macrophage (AM) populations in the lung^20^. We investigated whether CD11b^hi^ AM are preferentially infected following *Mtb* exposure and if their capacity to control the growth of clinical *Mtb* isolates of different lineages varied. We used an *in vivo* BCG vaccination and *Mtb-*infection model to examine which lung cells are *Mtb*-infected in the early stage of infection, and an *ex vivo* mycobacterial growth inhibition assay (MGIA) to assess the ability of AMs to restrict the growth of L2-B or L4 *Mtb*.

Mice were vaccinated with BCG and challenged six weeks later with fluorescently-labelled clinical strains of L2-B or L4 *Mtb* via aerosol **(Fig. 1A).** Lung tissues were collected at day 14 post-infection and analysed by flow cytometry to identify immune cell populations. Unbiased clustering revealed that while L2-B and L4 strains were detected across similar myeloid compartments, the population exhibiting the highest *Mtb* burden - defined by elevated TdTomato MFI - differed according to vaccination status. In unvaccinated mice, this signal was concentrated in cluster K10, which was subsequently characterised as predominantly Ly6G⁺Ly6C⁺, consistent with a neutrophil subset. In contrast, in BCG-vaccinated mice, peak TdTomato signal localised to cluster K17, characterised by CD11b^hi^CD64^hi^SiglecF^lo^ expression, consistent with a subset of AMs, suggesting a role for these cells in the early vaccinated lung response **(Fig. 1B).**

**Figure 1.**
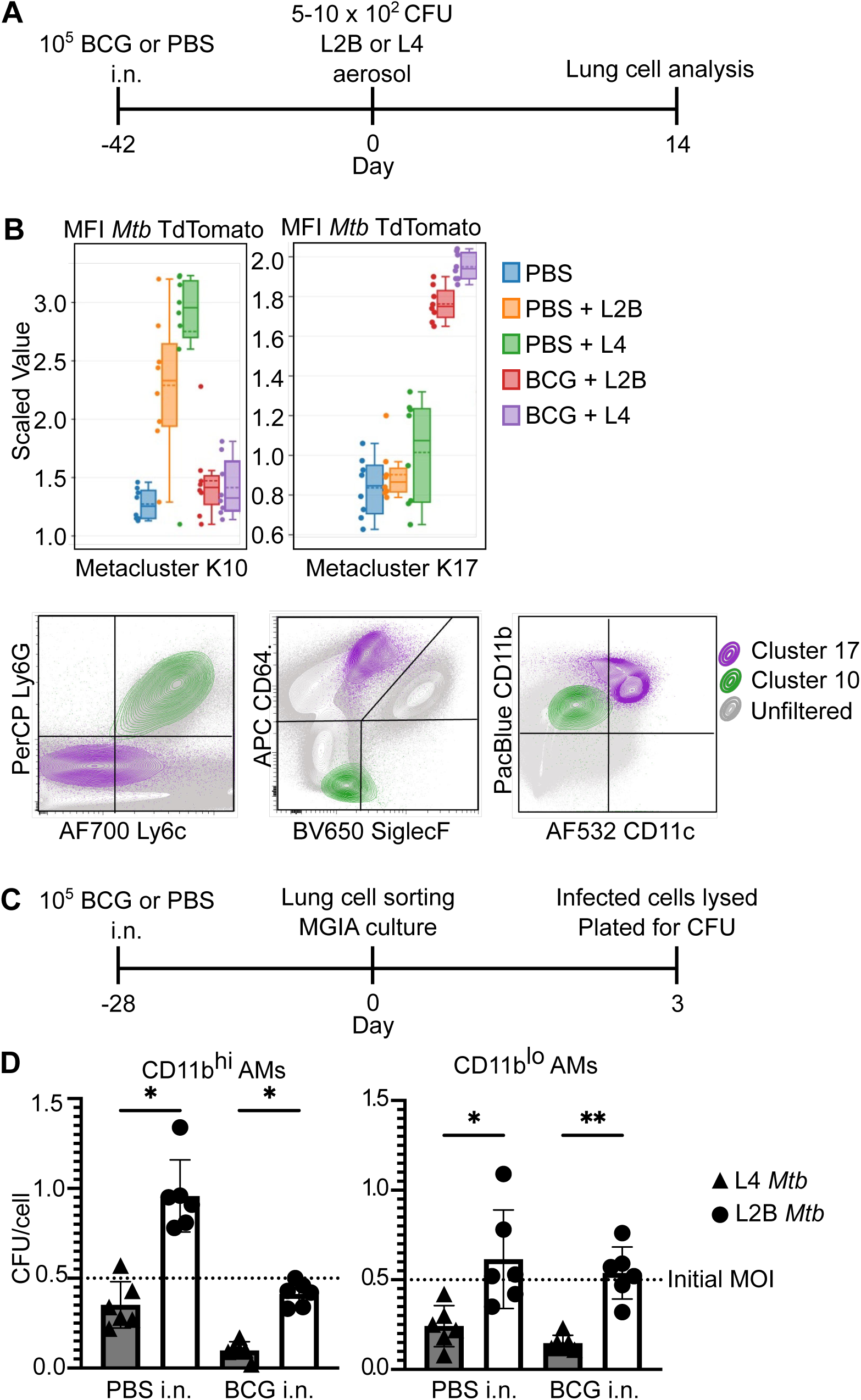
BCG-trained CD11b^hi^ alveolar macrophages restrict *Mtb* growth. A. Experimental design. Mice (n= 4m, 4f) were vaccinated with BCG and six weeks later challenged via aerosol with fluorescently labelled L2-B or L4 *Mtb* strains. Lungs were collected at days 1, 7, and 14 post-infection for flow cytometric analysis. B. FlowSOM analysis in OMIQ software provided unbiased metaclustering of lung immune cells, showing infection (defined as high MFI for TdTomato+ *Mtb*) was occurring in Ly6G+ Ly6C+ neutrophils (cluster K10) in unvaccinated mice and CD11b^hi^ CD64^hi^ SiglecF^lo^ alveolar macrophages (cluster K17) in BCG-vaccinated mice. C. Workflow for ex vivo infection assay. CD11b^hi^ and CD11b^lo^ alveolar macrophages were sorted from BCG-vaccinated lungs (n=9, 3 lungs pooled into 3 samples) and co-cultured with L2-B or L4 *Mtb* strains for 3 days to assess bacterial growth. D. Intracellular bacterial burden after 3-day co-culture, shown as CFU per cell for each macrophage subset and infecting strain. Plotted data represent 3 biological replicates from each of two experiments combined; boxes show median, whiskers indicate two standard deviations from the median.

To assess the functional capacity of CD11b^hi^ AMs, we isolated CD11b^hi^ and CD11b^lo^ AM subsets from BCG-vaccinated mouse lungs and infected them *ex vivo* with L2-B or L4 *Mtb* strains **(Fig. 1C).** After three days of co-culture, bacteria were quantified by CFU enumeration on agar. CD11b^hi^ AMs effectively restricted the intracellular growth of L4 *Mtb*, with marked reduction in CFU, consistent with active bacterial killing. In contrast, L2-B strains persisted at higher levels, indicating resistance to macrophage-mediated growth restriction **(Fig. 1D).** CD11b^lo^ AMs displayed limited capacity to restrict *Mtb* growth, regardless of bacterial lineage.

Together, these data show that in BCG-vaccinated mice, CD11b^hi^ AMs are the primary infected cell population in the lung during the first weeks of *Mtb* infection. *Ex vivo*, CD11b^hi^ AMs restricted *Mtb* growth more effectively than CD11b^lo^ cells, with reduced control of L2-B compared to L4.

### L2-B *Mtb* infection induces different cellular infiltration and interactions than L4 *Mtb* infection in BCG-vaccinated mice

Having established a key role for CD11b^hi^ AMs in BCG-vaccinated mice and their reduced control of L2-B *Mtb* strains, to interrogate the early evasive mechanisms of L2-B *Mtb in vivo*, we performed spatial transcriptomic analysis in the lungs of BCG-vaccinated mice at day 7 post-infection **(Fig. 2).** Mice were vaccinated intranasally with BCG and challenged six weeks later with L2-B or L4 *Mtb*. Lung tissues were formalin-fixed, paraffin-embedded, and sectioned onto 10x Genomics Visium slides for transcriptomic profiling.

**Figure 2.**
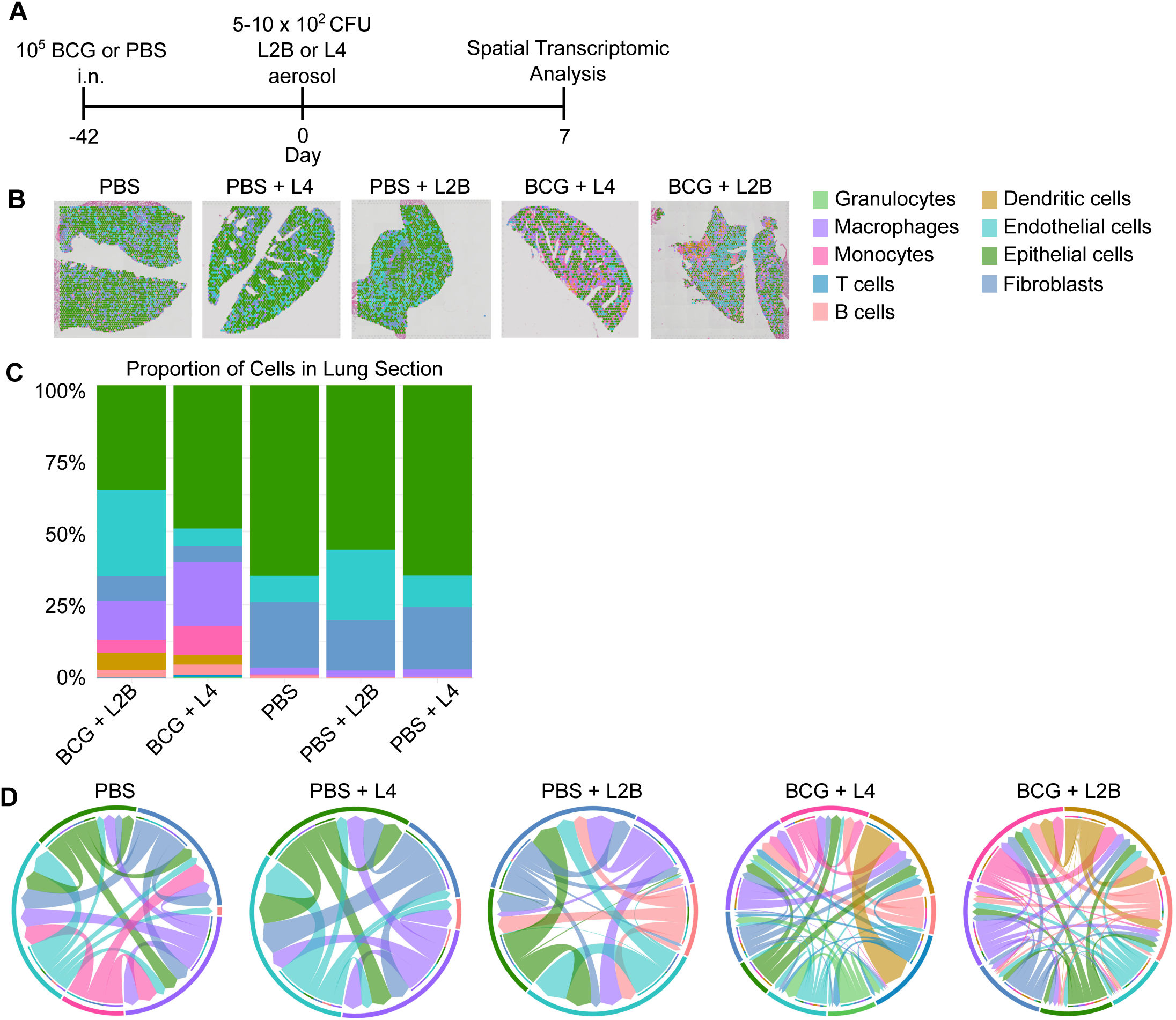
Spatial distribution and immune cell interactions in *Mtb*-infected lungs following BCG vaccination. A. Experimental design. Mice (n=2) were vaccinated intranasally with BCG and challenged six weeks later with L2-B or L4 *Mtb*. Lungs were collected at day 7 post-infection, formalin-fixed, paraffin-embedded, and analysed using 10x Visium spatial transcriptomics. B. Spatial plots showing dominant inferred cell type distribution across lung sections in each experimental group. C. Quantification of inferred immune cell representation across groups, based on refined spot-level cell type assignments from spatial transcriptomic data. D. CellChat analysis showing inferred cell–cell communication networks between major immune cell types in each group, based on refined spot-level cell type assignments.

Analysis of cell type composition across lung sections revealed increased immune cell infiltration in BCG-vaccinated groups compared to unvaccinated controls **(Fig. 2B,C).** Macrophages, monocytes, granulocytes, dendritic cells, and B cells were more abundant in vaccinated lungs, irrespective of the infecting *Mtb* strain. In contrast, unvaccinated lungs exhibited lower immune cell presence at this early time point.

To explore cell–cell communication, we applied CellChat analysis to infer intercellular signalling networks from spatial transcriptomic data **(Fig. 2D).** In both vaccinated and unvaccinated groups, L2-B-infected lungs showed prominent B cell interactions, whereas L4-infected lungs demonstrated more interactions involving granulocytes and T cells. These differences suggest variation in early immune network engagement between L2-B and L4 infections, particularly within the vaccinated lung environment.

### L2-B *Mtb* infection impairs innate immune activation and antimicrobial function

To investigate the molecular basis of the differential immune responses to L2-B and L4 *Mtb* observed in the lung, we performed differential gene expression (DEG) analysis using spatial transcriptomics **(Fig. 3).**

**Figure 3.**
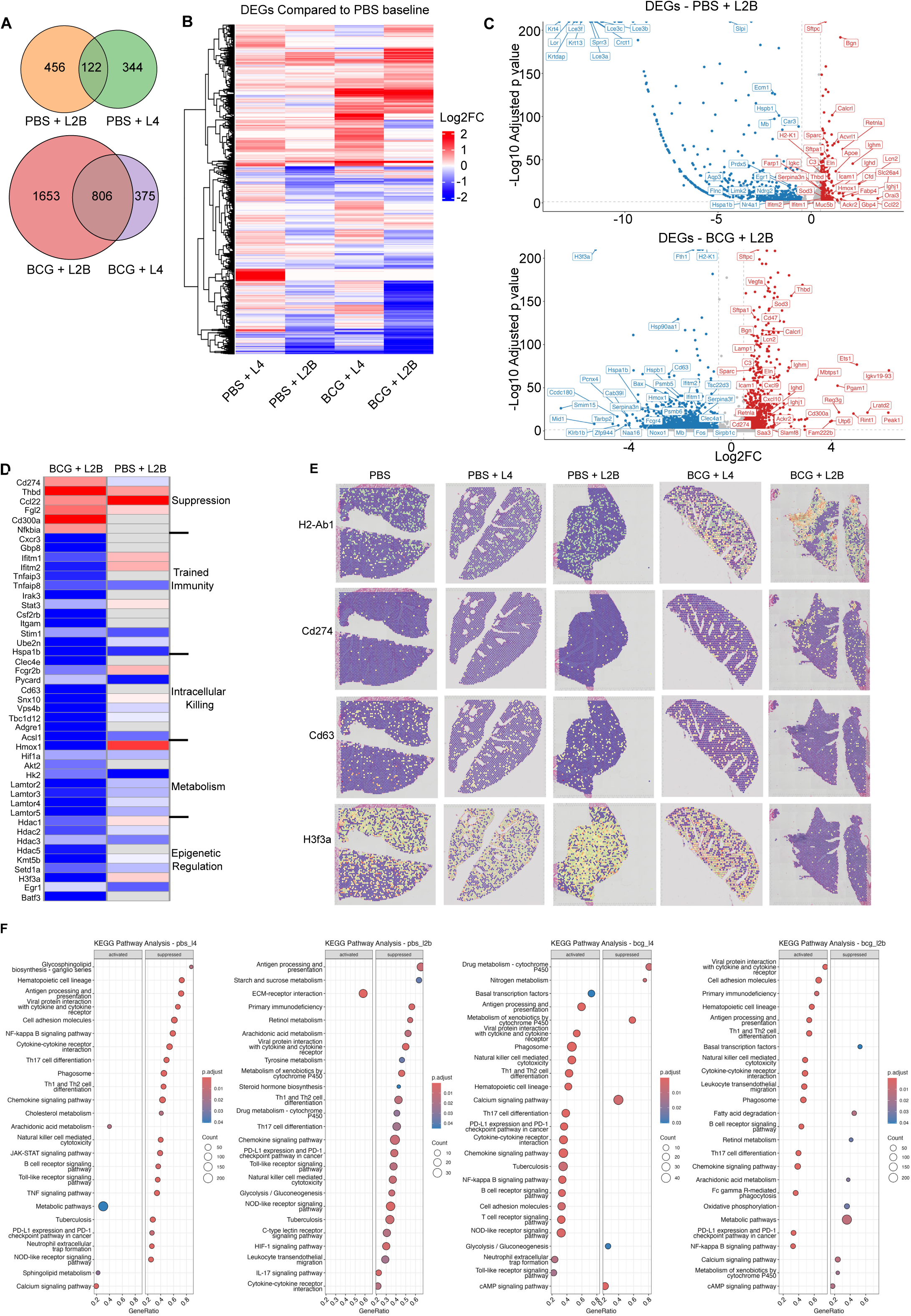
Differential gene expression analysis of L2-B- and L4-infected lungs using spatial transcriptomics. Mice (n=2) were vaccinated intranasally with BCG and challenged six weeks later with L2-B or L4 *Mtb*. Lungs were collected at day 7 post-infection, formalin-fixed, paraffin-embedded, and analysed using 10x Visium spatial transcriptomics. A. Venn diagrams showing the number of differentially expressed genes (DEGs) between L2-B- and L4-infected lungs in unvaccinated (PBS) and BCG-vaccinated groups. B. Heatmap showing global gene expression profiles across experimental groups. C. Volcano plot of DEGs comparing L2-B-and L4-infected lungs in BCG-vaccinated or unvaccinated (PBS) mice. D. Heatmap of immune-relevant DEGs grouped by functional category. E. Spatial expression maps displaying the distribution of selected immune-relevant genes in representative lung tissue sections. F. KEGG pathway enrichment analysis of L2-B- and L4-infected lungs in BCG-vaccinated or unvaccinated (PBS) mice. Only pathways relevant to immunological and host–pathogen processes are shown in the main figure for clarity; complete enrichment outputs are provided in Supplementary Tables 3 and 4.

Substantial numbers of DEGs were identified between L2-B and L4 infections in both groups, with a greater number of unique DEGs detected in BCG-vaccinated lungs **(Fig. 3A).** Global gene expression patterns showed widespread downregulation of transcripts in L2-B-infected lungs relative to L4, irrespective of vaccination status **(Fig. 3B).**

L2-B infection was associated with differential expression of genes that suggest impairment of an effective immune response **(Fig. 3C),** particularly in the context of BCG vaccination **(Fig. 3D).** Genes associated with trained immunity and innate immune memory, including *Cxcr3*, *Gbp8*, *Ifitm1/2*, *Tnfaip3*, *Irak3*, *Stat3*, and *Csf2rb*, were downregulated; notably *Itgam* (encoding CD11b), was suppressed, suggesting impaired myeloid cell recruitment or activation. Expression of antimicrobial genes critical for phagolysosome function and intracellular killing, such as *Clec4e*, *Fcgr2b*, *Pycard*, *Bnip3*, *Cd63*, and components of the endocytic trafficking machinery (*Snx10*, *Rab11b*, *Vps4b*, *Tbc1d12*, *Adgre*), were also suppressed.

L2-B infection was further associated with reduced expression of metabolic regulators such as *Hif1a*, *Acsl1*, *Hmox1*, *Nqo1*, *Aldh3a1*, *Akt2*, *Hk2*, and multiple components of the *Lamtor* complex, implicating compromised immune cell bioenergetics. Additionally, key regulators of chromatin accessibility and transcriptional memory were downregulated, including: *Hdac1-3*, *Hdac5*, *Kmt5b*, *Setd1a*, *H3f3a*, *Egr1*, and *Batf3*, suggesting that epigenetic reprogramming may underlie the observed immune suppression. By contrast, genes with immunoregulatory or suppressive potential, such as *Cd274* (PD-L1), *Thbd*, *Ccl22*, *Fgl2*, *Cd300a*, and *Nfkbia*, were upregulated. Together, these transcriptional changes point to lineage-specific immune evasion strategies employed by L2-B that may limit effective BCG-trained innate responses.

Spatial transcriptomic profiling of lung tissue sections revealed that although genes such as *H2-Ab1* (MHCII) remained expressed in L2-B-infected lungs - indicative of ongoing immune cell presence - this was accompanied by suppression of genes essential for an effective antimicrobial response. These transcriptional alterations were not restricted to specific tissue regions but were diffusely distributed throughout the lung **(Fig. 3E).** Immunoregulatory genes such as *Cd274* (PD-L1) were notably upregulated in L2-B, while *Cd63*, involved in the maturation of phagosomes, was downregulated. Of particular interest, expression of *H3f3a*, a histone variant implicated in chromatin remodeling and trained innate immunity, was markedly reduced in L2-B-infected lungs, exclusively in BCG-vaccinated mice.

Pathway analysis further illustrated these transcriptional differences. Kyoto Encyclopedia of Genes and Genomes (KEGG) enrichment was used to assess broader pathway-level activation and suppression across groups. In unvaccinated lungs, while key immune pathways were predominantly suppressed after *Mtb* infection with either lineage, the response to L4 involved strong activation of metabolic pathways, while L2-B-infection resulted in unique suppression of pathways including HIF-1 signalling and multiple metabolism-associated pathways including glycolysis/gluconeogenesis (Fig. 3F). BCG vaccination activated the majority of these pathways, however suppression of basal transcription factors, fatty acid degradation and metabolism-related pathways persisted in L2-B-infected lungs.

To further define which host-response processes were uniquely suppressed in L2-B relative to L4, we performed a targeted supplementary GO analysis focused on biologically relevant immune, antimicrobial, metabolic, and transcriptional pathways (**Supplementary Fig. 1**). In unvaccinated mice, this highlighted suppression of pathways related to defence responses to bacteria, phagocytosis, and regulation of innate immune responses. In BCG-vaccinated mice, L2-B infection was associated with suppression of host-response terms linked to mitochondrial function, oxidative phosphorylation, fatty acid and glucose metabolism, RNA processing including rRNA metabolic pathways, and selected DNA-associated processes.

These analyses reveal that L2-B infection disrupts multiple layers of the host immune response - including microbial killing, immune signalling, metabolism, and transcriptional activity - despite robust immune cell infiltration and evidence of antigen presentation, potentially contributing to its enhanced capacity to evade BCG-mediated immune control.

### L2-B and L4 infections trigger distinct transcriptional regulatory networks

To investigate upstream transcriptional regulation contributing to the differential immune responses observed in L2-B- versus L4-infected lungs, we performed GENIE3-based regulatory network inference on spatial transcriptomic data. Heatmap analysis of the top 20 most variable transcription factors (TFs) relative to PBS control revealed a global reduction in inferred regulatory strength in L2-B-infected lungs, evident in both unvaccinated and BCG-vaccinated groups **(Fig. 4A).** By contrast, L4-infected lungs exhibited stronger inferred TF regulatory activity.

**Figure 4.**
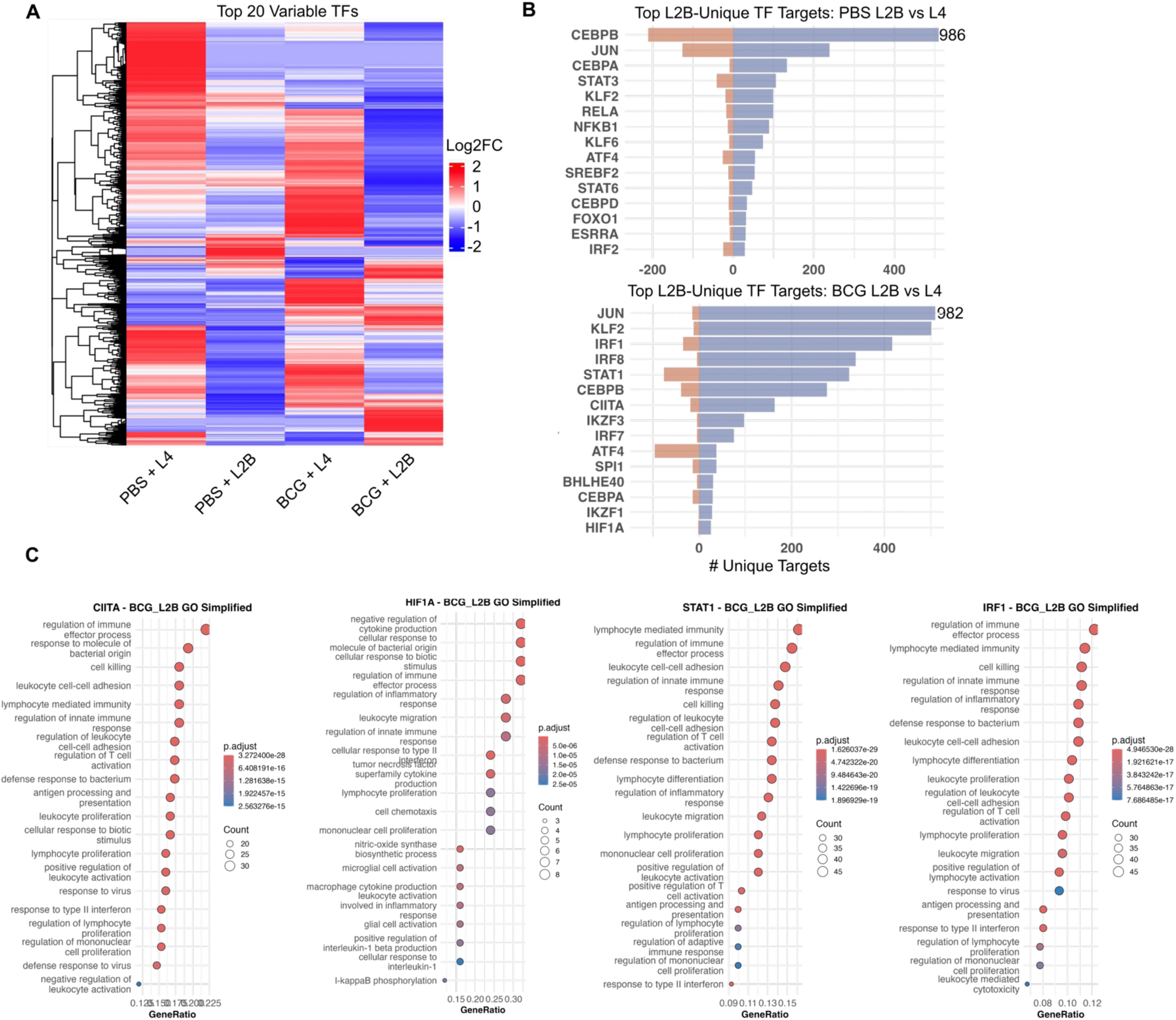
Transcription factor network inference reveals strain-specific inferred regulatory programmes in L2-B- and L4-infected lungs. Mice (n=2) were vaccinated intranasally with BCG and challenged six weeks later with L2-B or L4 *Mtb*. Lungs were collected at day 7 post-infection, formalin-fixed, paraffin-embedded, and analysed using 10x Visium spatial transcriptomics. A. Heatmap showing the top 20 most variable transcription factors (TF) across experimental groups, based on average inferred regulatory strength from GENIE3 networks and displayed relative to PBS control. B. Bar plot showing the number of unique inferred target genes regulated by individual TFs in BCG-vaccinated or unvaccinated mice, infected with L2-B or L4. C. Gene Ontology (GO) enrichment analysis of target genes uniquely regulated by selected TFs in BCG-vaccinated L2-B-infected lungs, defined as targets present in L2-B but not L4 inferred regulatory networks. Regulators shown (CIITA, STAT1, IRF1, and HIF1A) were chosen from the inferred network analysis on the basis of broad regulatory control and biological relevance. Simplified GO terms are displayed with enrichment categories and associated gene counts.

We next examined inferred regulatory strength across transcriptional regulators known to influence immune responses. In BCG-vaccinated mice, L2-B infection was associated with reduced regulatory strength of several transcriptional regulators known to influence macrophage activation, trained immunity, and metabolic programming, including CHD1, MAFB, FOS, IRF8, and TRAF6 **(Supplementary Fig. 2)**. Conversely, TFs such as RELA, STAT3, FOXO1, PPARG, and CEBPB showed increased regulatory strength in L2-B compared to L4, potentially affecting inflammatory modulation, macrophage polarisation, and stress adaptation. In unvaccinated mice, L2-B infection showed increased regulatory strength in TFs including BATF, IRF1, IRF4, XBP1, and ATF4, several of which have been linked to ER stress, macrophage adaptation, and effector programming. Meanwhile, TFs such as IRF9, SREBF2, FOXO1, and STAT6 were reduced in activity compared to L4, suggesting that L2-B infection interferes with transcriptional programs associated with type I interferon signalling, lipid metabolism, cellular stress adaptation, and Th2 or regulatory immune responses.

Notably, only a small subset of regulators (e.g., MAFB, EZH2, STAT5B, CHD1, ZBTB32, and PPARGC1B) exhibited consistently lower inferred regulatory strength in L2-B-infected lungs compared to L4, regardless of BCG vaccination status. These TFs are broadly involved in epigenetic regulation, macrophage differentiation, and immune-metabolic programming. Conversely, several TFs - including BATF, DDIT3, NFKB2, STAT4, and TFE3 - were uniquely upregulated in L2-B infection across both vaccinated and unvaccinated mice, implicated in stress responses, pro-inflammatory signalling, and lymphoid cell activation. These shared regulatory patterns may reflect a core feature of L2-B-associated host reprogramming across immune contexts, in which transcriptional programmes supporting antimicrobial capacity and immune memory are diminished, while other TF changes appear to be vaccination-context dependent.

We next examined the specific downstream gene networks associated with individual TFs in each group to better understand the functional implications of this strain-specific control. We assessed the number of unique inferred target genes associated with individual TFs in L2-B-versus L4-infected lungs. In both PBS-treated and BCG-vaccinated mice, a small number of TFs were associated with a large number of unique inferred target genes in L2-B-infected lungs compared to L4-infected lungs **(Fig. 4B).** These findings suggest that, although global inferred TF regulatory strength is diminished in L2-B infection, specific TFs retain or gain influence over distinct predicted gene networks, which may drive strain-specific immune modulation despite overall suppression of protective responses.

To assess the functional implications of these strain-specific regulatory networks, we performed GO enrichment analysis on the unique inferred target gene sets associated with selected TFs in BCG-vaccinated L2-B-infected lungs, defined as genes uniquely regulated in L2-B compared to L4. GO enrichment results were subsequently simplified by grouping and filtering related terms to reduce redundancy and highlight interpretable TF-specific regulatory themes. Predicted target genes of CIITA were enriched for pathways related to regulation of immune and defence responses, predicted STAT1 targets were associated with cell-mediated immunity and antimicrobial responses, while predicted IRF1 targets included genes involved in leukocyte migration and effector functions **(Fig. 4C).** Interestingly, predicted HIF1A target genes were enriched for pathways linked to nitric oxide metabolism, cytokine production, and hypoxia responses, consistent with metabolic alterations observed in L2-B infection.

These data demonstrate that L2-B infection is associated with distinct inferred transcriptional regulatory networks, with specific TFs linked to divergent predicted gene sets. This strain-specific reprogramming of host responses may contribute to the differential immune outcomes observed between L2-B and L4 infections.

### Tuberculosis household contacts exposed to L2 strains show higher rates of *Mtb* **infection**

To determine if the early responses identified in the mouse lung were mirrored in humans, we analysed peripheral blood from individuals with household exposure to tuberculosis patients infected with either L2 or L4 strains. The INFECT cohort study in Indonesia examined 1347 household contacts of sputum smear-positive tuberculosis patients^21^. *Mtb* infection, as detected by interferon gamma (IFN-γ) release assay (IGRA), was measured in household contacts initially and again after three months for those who tested negative at baseline. Household contacts who remained IGRA-negative at three months were defined as ‘early clearers’^22^, while those who became IGRA-positive were termed ‘IGRA converters’. IGRA conversion among contacts was associated with higher exposure (e.g. closer sleeping proximity to or higher sputum bacterial load of the index patient), while the presence of a BCG scar correlated with reduced risk of IGRA conversion (aRR 0.56; 95% CI, 0.40 - 0.77).

To examine the effect of *Mtb* genotype on the risk of infection, we linked IGRA conversion among household contacts with the index patient’s *Mtb* genotype, based on whole genome sequencing (WGS) of cultured sputum *Mtb* isolates^6^. We excluded individuals who had positive baseline IGRA results, as a significant proportion could be from prior infection with an unknown *Mtb* isolate rather than recent infection with the index patient’s strain.

Among 375 baseline IGRA-negative contacts from households with an available *Mtb* genotype, no significant differences were found in clinical characteristics between those exposed to L2 (L2-B and non-Beijing L2, n=114) and L4 (n=261) *Mtb* strains (**Table 1**). However, IGRA conversion appeared more common among household contacts exposed to L2 compared to L4 strains (34 vs 25% with the standard IGRA cut-off *P* value = 0.06; 27 vs 15% with a stricter IGRA cut-off, *P* value = 0.03). Moreover, among those with IGRA conversion, quantitative TB-specific responses were significantly higher among those exposed to L2 strains compared to L4 (**Fig. 5A**). When restricted to contacts exposed to the L2-B sublineage, IGRA conversion was even more frequent, and quantitative IGRA results were higher compared to non-Beijing L2 (**Supplementary Table 1**).

**Figure 5.**
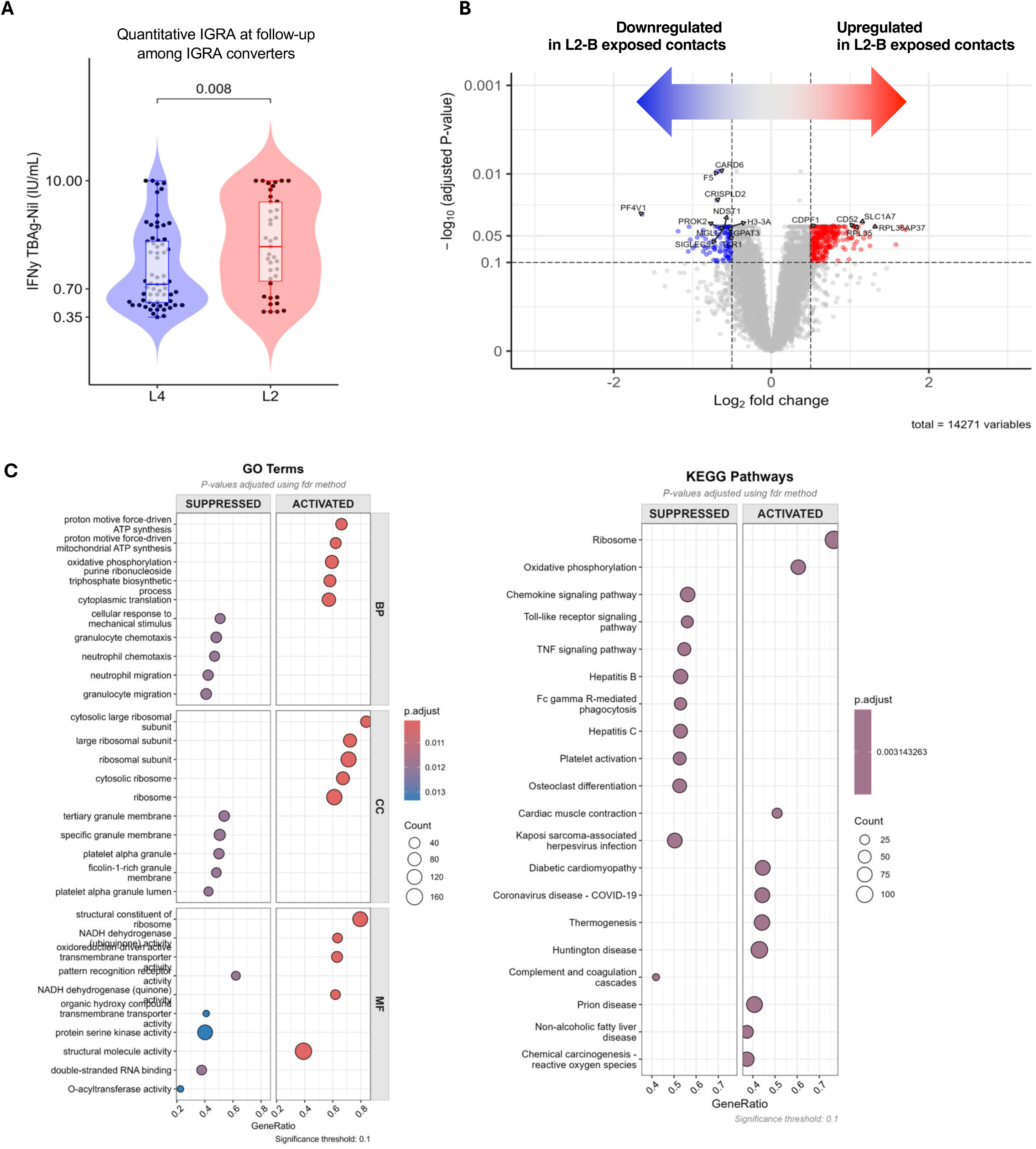
Early transcriptional responses differ in household contacts exposed to L2 versus L4 *Mtb*. A. IGRA converters exposed to L2 *Mtb* have a higher quantitative IGRA response compared to IGRA converters who were exposed to L4 *Mtb*. B. Volcano plot showing differential gene expression between household contacts exposed to L2-B (n=15) versus L4 (n=24) *Mtb* strains at baseline, with 1488 significantly differentially expressed genes after multiple testing correction (adjusted P value [FDR] < 0.05). Blue points represent genes downregulated in L2-B-exposed contacts, red points represent upregulated genes. Analysis was adjusted for age and IGRA status at follow-up to isolate strain-specific effects. C. Gene Ontology (GO) biological process enrichment analysis of differentially expressed genes. Dot plot shows suppressed and activated pathways in L2-B-exposed contacts compared to L4-exposed contacts. Dot size represents gene count and colour intensity represents adjusted P value. D. KEGG pathway enrichment analysis showing pathway-level differences between exposure groups.

**Table 1.**
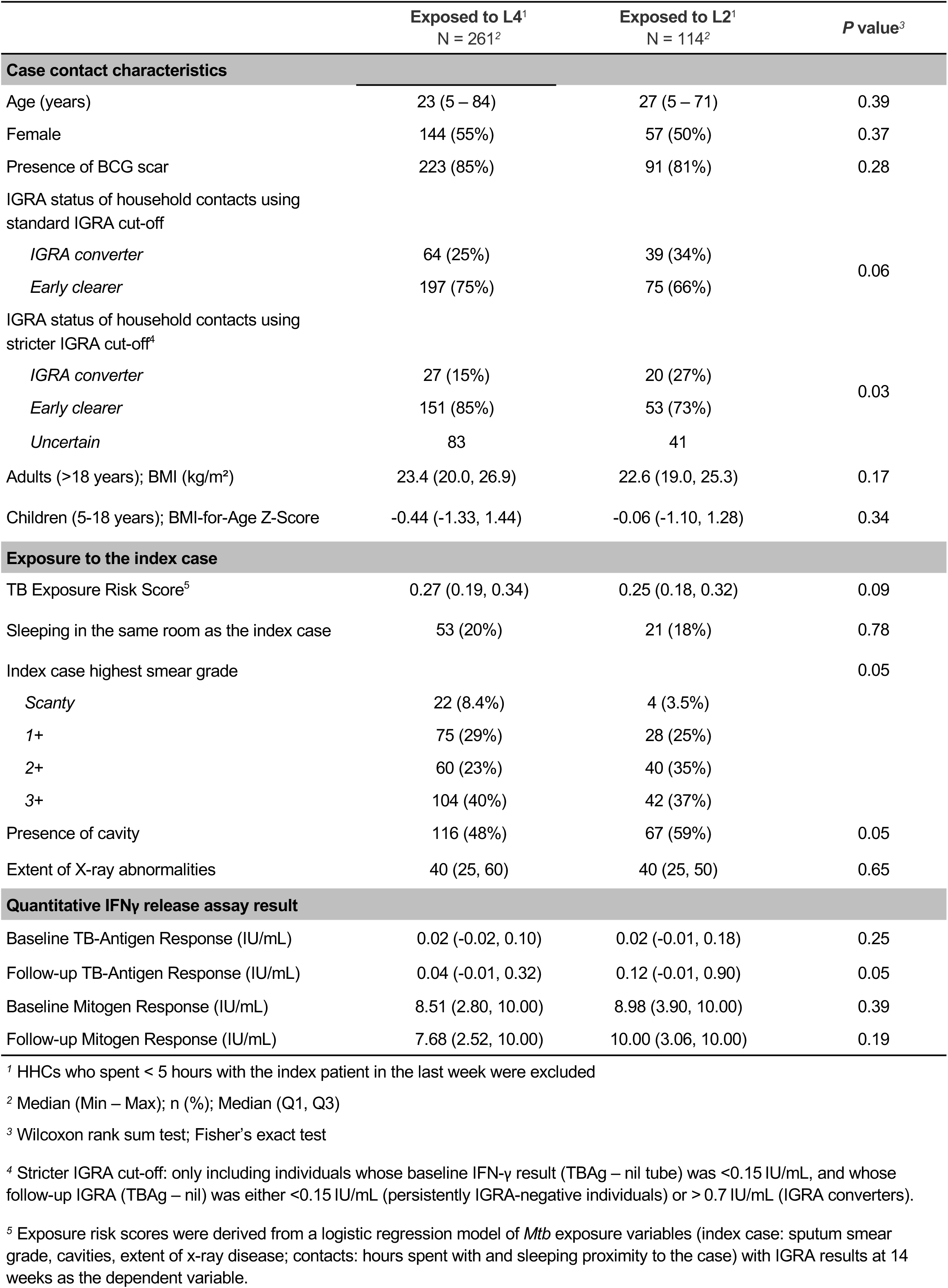
Household contacts characteristics stratified by *Mtb* lineage of index patients.

|  | Exposed to L4<sup>1</sup><br>N = 261<sup>2</sup> | Exposed to L2<sup>1</sup><br>N = 114<sup>2</sup> | P value<sup>3</sup> |
| --- | --- | --- | --- |
| **Case contact characteristics** |  |  |  |
| Age (years) | 23 (5 – 84) | 27 (5 – 71) | 0.39 |
| Female | 144 (55%) | 57 (50%) | 0.37 |
| Presence of BCG scar | 223 (85%) | 91 (81%) | 0.28 |
| IGRA status of household contacts using standard IGRA cut-off |  |  |  |
| *IGRA converter* | 64 (25%) | 39 (34%) | 0.06 |
| *Early clearer* | 197 (75%) | 75 (66%) |  |
| IGRA status of household contacts using stricter IGRA cut-off<sup>4</sup> |  |  |  |
| *IGRA converter* | 27 (15%) | 20 (27%) | 0.03 |
| *Early clearer* | 151 (85%) | 53 (73%) |  |
| *Uncertain* | 83 | 41 |  |
| Adults (>18 years); BMI (kg/m<sup>2</sup>) | 23.4 (20.0, 26.9) | 22.6 (19.0, 25.3) | 0.17 |
| Children (5-18 years); BMI-for-Age Z-Score | -0.44 (-1.33, 1.44) | -0.06 (-1.10, 1.28) | 0.34 |
| **Exposure to the index case** |  |  |  |
| TB Exposure Risk Score<sup>5</sup> | 0.27 (0.19, 0.34) | 0.25 (0.18, 0.32) | 0.09 |
| Sleeping in the same room as the index case | 53 (20%) | 21 (18%) | 0.78 |
| Index case highest smear grade |  |  | 0.05 |
| *Scanty* | 22 (8.4%) | 4 (3.5%) |  |
| 1+ | 75 (29%) | 28 (25%) |  |
| 2+ | 60 (23%) | 40 (35%) |  |
| 3+ | 104 (40%) | 42 (37%) |  |
| Presence of cavity | 116 (48%) | 67 (59%) | 0.05 |
| Extent of X-ray abnormalities | 40 (25, 60) | 40 (25, 50) | 0.65 |
| **Quantitative IFNγ release assay result** |  |  |  |
| Baseline TB-Antigen Response (IU/mL) | 0.02 (-0.02, 0.10) | 0.02 (-0.01, 0.18) | 0.25 |
| Follow-up TB-Antigen Response (IU/mL) | 0.04 (-0.01, 0.32) | 0.12 (-0.01, 0.90) | 0.05 |
| Baseline Mitogen Response (IU/mL) | 8.51 (2.80, 10.00) | 8.98 (3.90, 10.00) | 0.39 |
| Follow-up Mitogen Response (IU/mL) | 7.68 (2.52, 10.00) | 10.00 (3.06, 10.00) | 0.19 |

<sup>1</sup> HHCs who spent < 5 hours with the index patient in the last week were excluded<sup>2</sup> Median (Min – Max); n (%); Median (Q1, Q3)<sup>3</sup> Wilcoxon rank sum test; Fisher's exact test<sup>4</sup> Stricter IGRA cut-off: only including individuals whose baseline IFN-γ result (TBAg – nil tube) was <0.15 IU/mL, and whose follow-up IGRA (TBAg – nil) was either <0.15 IU/mL (persistently IGRA-negative individuals) or > 0.7 IU/mL (IGRA converters).<sup>5</sup> Exposure risk scores were derived from a logistic regression model of *Mtb* exposure variables (index case: sputum smear grade, cavities, extent of x-ray disease; contacts: hours spent with and sleeping proximity to the case) with IGRA results at 14 weeks as the dependent variable.

### Exposure to L2-B strains is associated with suppression of pathways related to innate immune responses

To investigate differences in the peripheral blood immune response, we compared whole blood transcriptomic profiles from household contacts exposed to either L2-B or L4 strains using standard IGRA cut-off classification. At baseline, when all individuals were IGRA-negative, whole blood RNA sequencing was compared between 15 L2-exposed contacts (all exposed to the L2-B sublineage) and 24 L4-exposed contacts. We found that 1488 genes were differentially expressed in contacts exposed to L2-B versus those exposed to L4, after correction for multiple testing and adjusting for age and IGRA status at follow-up (**Fig. 5B**).

Gene set enrichment showed suppression of pathways related to innate immune responses and activation of metabolic pathways in contacts exposed to L2-B. The GO biological processes (BP) terms that were suppressed in contacts exposed to L2-B were granulocyte and neutrophil chemotaxis and migration, while the activated terms were related to ATP production and oxidative phosphorylation (**Fig. 5C**). In the KEGG pathway analysis, again, pathways related to the innate immune response such as chemokine signalling pathways, Toll-like receptor signalling pathways, TNF signalling pathway and Fc-gamma receptor mediated phagocytosis were all suppressed in contacts exposed to L2-B *Mtb*, while oxidative phosphorylation was activated (**Fig. 5D**).

Since L2 was associated with IGRA conversion and quantitative IFN-γ production at month 3, we next looked at gene expression comparing contacts whose IGRA converted after exposure to L2-B (n=3) and L4 (n=5) strains (**Fig. 6A**). Even within this small subgroup, 283 genes were differentially expressed, after adjustment for age and correction for multiple testing (**Fig. 6B**).

**Figure 6.**
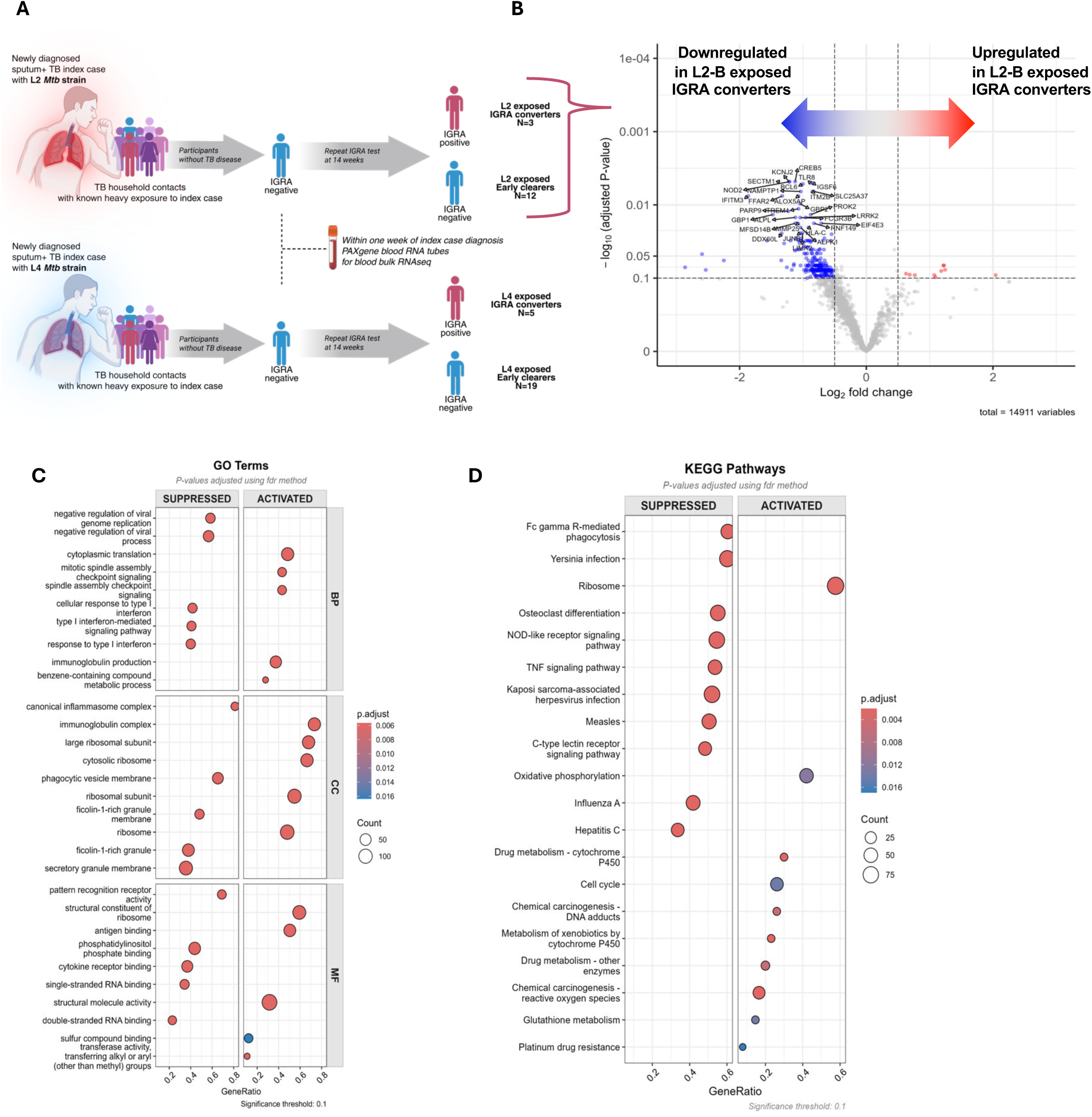
Strain-specific transcriptional differences in IGRA converters exposed to L2-B versus L4 *Mtb*. A. Study schematic showing household contacts exposed to either L2-B or L4 *Mtb* strains, with contacts classified as IGRA converters (red pathway) or early clearers (blue pathway) based on interferon gamma release assay results at 3-month follow-up. Transcriptomic analysis was performed on baseline blood samples from IGRA-negative contacts. B. Volcano plot showing differential gene expression in IGRA converters exposed to L2-B versus L4 strains (n=3 L2-B, n=5 L4). Blue arrows indicate downregulated genes and red arrows indicate upregulated genes in L2-B-exposed contacts. Significance threshold set at adjusted P value (FDR) < 0.1. A total of 283 genes were differentially expressed. C. Gene Ontology (GO) biological process terms enrichment analysis for IGRA converters. Dot plot showing suppressed and activated pathways in L2-B-exposed contacts compared to L4-exposed contacts. Dot size represents gene count and colour intensity represents adjusted P value significance. D. KEGG pathway enrichment analysis for IGRA converters showing suppressed and activated pathways.

Surprisingly, 95% of differentially expressed genes were downregulated upon exposure to L2-B strains in people whose IGRA later converted. In IGRA converters exposed to L2-B, baseline whole blood transcriptomics (when IGRAs were still negative) showed profound suppression of key immune pathways, including the type 1 IFN-mediated signalling pathway, cellular response to type 1 IFN, and pattern recognition receptor activity (**Fig. 6C**), as well as Fc-gamma receptor mediated phagocytosis, TNF signalling pathway, and NOD-like receptor signalling pathways (**Fig. 6D**). Interestingly, the pathways related to immunoglobulin production, immunoglobulin complex, and oxidative phosphorylation were activated. This suggests that early exposure to L2-B *Mtb* in individuals who later fail to control the infection is characterised by broad suppression of innate and cellular adaptive immune responses with activation of antibody production and oxidative phosphorylation very early after infection.

### Divergent transcriptional signatures underpin vaccine escape versus clearance

Comparison of mouse and human datasets identified overlapping transcriptional features between individuals who converted to IGRA positive following L2-B exposure and BCG-vaccinated mice infected with L2-B *Mtb*. These shared gene sets were enriched for pathways related to B cell and immunoglobulin responses, alongside reduced representation of innate immune signalling pathways. By contrast, early clearers displayed a distinct transcriptional profile, with enrichment of pathways linked to regulation of trained innate immunity and cellular metabolism, which were not observed in the converter-associated or L2-B mouse gene sets (**Supplementary Fig. 3**). Together, these patterns suggest candidate transcriptional programs associated with vaccine escape versus early clearance.

## Discussion

Despite widespread administration, BCG vaccination offers variable protection against tuberculosis. Evidence from household contact studies indicates a modest reduction in the risk of *Mycobacterium tuberculosis* infection, as measured by interferon-γ release assay conversion, among vaccinated children following exposure^23^. Protection against tuberculosis disease has been observed primarily in early childhood and is reduced in older age groups^24^. The mechanisms underlying this limited and heterogeneous protection remain poorly understood. In this study, we investigate how BCG-induced immunity is circumvented by modern L2-B strains using geographically matched clinical isolates of L2-B and L4 *Mtb*.

Trained immunity is gaining recognition as a critical component of BCG-mediated protection. BCG vaccination induces functional reprogramming in innate cells, particularly monocytes and macrophages, through histone modifications and metabolic shifts toward aerobic glycolysis, driven via the Akt–mTOR–HIF-1α axis^11–13^. Lung-resident AMs have been shown to acquire memory-like features following systemic BCG vaccination^15^ and expand and adopt a CD11b^hi^ phenotype in response to intranasal BCG vaccination^20^. These trained AMs display rapid and enhanced responses to secondary *Mtb* exposure, and emerging evidence suggests they are key players in early containment of infection^16^.

In our study, CD11b^hi^ AMs were the dominant infected population in BCG-vaccinated mice, early after exposure to either L2-B or L4 *Mtb*. BCG-trained AMs restricted growth of L4 *Mtb* but failed to control L2-B growth, suggesting that L2-B *Mtb* strains can evade BCG-mediated AM training. This implies that hypervirulent *Mtb* strains possess mechanisms to limit the efficacy of trained immunity, and has particular public health relevance, as L2-B strains are highly prevalent in Asia and are associated with resistance to protection afforded by the BCG vaccine^6^. This suggests that the global variation in the efficacy of BCG may, in part, be explained by lineage-specific strategies of immune evasion.

Spatial transcriptomic profiling of infected lung tissue revealed transcriptional and regulatory reprogramming, with widespread suppression of pathways essential for macrophage activation and trained innate immunity, providing mechanistic insight into why BCG-trained immune cells fail to control L2-B. In contrast to the immune activation seen during L4 infection, L2-B-infected lungs showed broad downregulation of antimicrobial defence genes and phagosome-lysosome biology, and humans exposed to L2 *Mtb* showed suppression of Fc gamma R-mediated phagocytosis. Consistent with this, *Cd63*, encoding a tetraspanin required for late endosomal trafficking and phagolysosomal maturation, was specifically reduced in BCG-vaccinated mice infected with L2-B, indicating ineffective maturation of phagosomes despite the presence of trained macrophages. In addition, metabolic regulators - including *Hif1a*, *Hmox1*, *Akt2*, and *Lamtor* components - were reduced in infected mice, demonstrating the *in vivo* undermining of metabolic processes essential for functional trained immunity^12,13^.

A significant finding of this study is that expression of the histone variant *H3f3a* (H3.3) was profoundly silenced, together with its chaperone *Asf1a*, alongside downregulation of chromatin remodellers that interplay to maintain transcriptionally permissive chromatin^25,26^. This effect was evident specifically in BCG-vaccinated mice infected with L2-B, but not in unvaccinated mice or with L4, suggesting that this mechanism is not a default virulence trait but is specifically deployed under the selective pressure of BCG-induced trained immunity.

Consistent with this, we also observed reduced H3.3 expression in peripheral blood of humans exposed to L2-B compared with L4. Given that the majority of human contacts in this cohort had prior BCG vaccination, the concordant suppression upon exposure to L2-B highlights that this escape mechanism is not limited to experimental models but is deployed under real-world vaccine pressure. Loss of H3.3 has been shown to lead to reduced haematopoietic stem cell (HSC) numbers and biased myelopoiesis, and is required for the maintenance of adult HSC stemness^27^. The importance of BCG mediated training of HSCs for protective innate immunity to tuberculosis has been described^15^, and *Mtb* shown to be capable of reprogramming HSCs^28^.

The striking, vaccine-restricted suppression of H3.3 expression observed in our study coincided with reduced glycolytic and innate immune responses *in vivo*, and was functionally reflected in the impaired ability of BCG-trained macrophages to control L2-B growth in a MGIA. The speed (within one week after infection in mice, and early after exposure to an infectious contact in humans) and specificity of *H3f3a* loss suggest an active process by which L2-B manipulates host chromatin. Given that H3.3 is continuously turned over and relies on active deposition machinery (such as *Asf1a*), interference with these pathways could rapidly extinguish *H3f3a* and associated chromatin accessibility. Thus, L2-B may escape BCG-induced trained immunity through early, targeted disruption of histone variant supply and enhancer maintenance, allowing L2-B to actively disarm the epigenetic and metabolic programs that underpin BCG-induced trained immunity.

This model is supported by existing evidence that *Mtb* can manipulate host chromatin and innate immune signalling through secreted effectors. For example, *Mtb* encodes a secreted histone methyltransferase, Rv1988, which can translocate into the host nucleus and methylate histone H3 to repress immune gene transcription^29^. BCG-trained macrophages will likely have chromatin open at inflammatory enhancers (via H3.3, H3K4me3, H3K27ac), this “poised” state making them more responsive but potentially more vulnerable to manipulation. L2-B may only be able to target pathways that are active; in unvaccinated lungs, where H3.3 turnover is lower and enhancers are relatively closed, the bacterium may not target this pathway. Other mechanisms include the production of a phenolic glycolipid by some L2-B isolates, which inhibits TLR signalling and dampens early macrophage release of TNF, IL-6, and IL-12^4^, as well as the capacity of virulent strains to skew host responses toward a detrimental type I IFN signature while suppressing protective IL-1 production^30,31^. In addition, the *Mtb* ESX-1 secretion system delivers bacterial effectors such as ESAT-6 into host cells, disrupting phagosomal membranes and altering downstream immune signalling^32^.

Transcription factor (TF) network analysis further clarified how these transcriptional changes may be orchestrated. L2-B infection was associated with reduced regulatory strength of CHD1, MAFB, FOS, IRF8, and TRAF6. These TFs are involved in macrophage activation and trained innate immunity, with CHD1 acting as a chromatin remodeller^33^, and IRF8 known to shape enhancer activity and chromatin accessibility during cell differentiation^34^. Suppression of these regulators therefore provides a plausible mechanism by which L2-B undermines both transcriptional activation and the epigenetic reprogramming required for durable trained responses. In contrast, a smaller set of TFs (BATF, DDIT3, STAT4, TFE3) was selectively upregulated, but these TFs were linked primarily to stress adaptation, cytokine signalling, or alternative macrophage polarisation rather than direct antimicrobial defence. This pattern suggests that L2-B rewires TF hierarchies away from protective programs and toward pathways that favour immune evasion and cellular tolerance.

In support of this, several TFs, including *IRF1*, *STAT1*, *CEBPB*, *CIITA*, *IRF8* and *HIF1A*, had strikingly large numbers of unique target genes in L2-B-infected lungs following BCG vaccination (Fig. 4), but not in response to L4 *Mtb*. These TFs are each known to regulate core aspects of antimycobacterial immunity. For instance, *IRF1* and *STAT1* are central to IFN-γ and IL-1 responses^35^, *HIF1A* controls metabolic training and IL-1β production^12^, *CEBPB* shapes macrophage activation and cytokine production^36^, and *CIITA* regulates MHC-II antigen presentation^37^. Previous reports show *STAT1* and *IRF1* can be exploited to promote type I IFN responses that drive TB progression^35^. These TFs may be re-engaged by L2-B to activate alternative, maladaptive programs rather than protective ones.

Interestingly, L2-B infection was also associated with upregulation of immunosuppressive or regulatory genes, including *Cd274* (PD-L1), *Thbd* (thrombomodulin), and *Ccl22* (Fig. 3). PD-L1 is a known immune checkpoint that inhibits T cell and macrophage responses during *Mtb* infection^38^, while CCL22 promotes recruitment of regulatory T cells or Th2 polarisation^39^. L2-B infection in the mouse lung was associated with an increase in cell-cell communication involving B cells, with increased immunoglobulin pathway activity in the human cohort being associated with those who went on to become IGRA converters. Consistent with this, cross-species comparison of DEGs showed that both converters and L2-B-infected mice shared a maladaptive signature characterised by B cell and immunoglobulin pathway upregulation and suppression of innate immune responses, whereas early clearers displayed a distinct protective program marked by enhanced oxidative phosphorylation and translational activity. In keeping with this, a study comparing the course of Beijing-1585 (L2-B) *Mtb* infection with H37Rv (L4) *Mtb* infection in unvaccinated mice found that mice infected with L2-B had an increased percentage of B cells in the lung at day 14 after intratracheal infection, concomitant with increased production of the canonical Th2 cytokine, IL-4.^40^ These factors may suggest a suboptimal immune response is established early after infection. These observations also align with prior work showing increased Treg recruitment during L2-B *Mtb* infection^7,8^ and extend the timeline to suggest that suppression begins well before adaptive immunity is established.

A striking aspect of our human study is that individuals who later became IGRA converters already showed a perturbed immune response in their baseline RNA-seq profiles, taken very early after exposure to L2-B while they were still IGRA-negative. These transcriptional defects appearing before systemic immune conversion provide a rare window into the earliest phase of human infection, suggesting that failure of protective immunity may be programmed very early after exposure. The observation that early transcriptomic signatures predicted subsequent IGRA conversion raises the possibility that peripheral blood sampling could provide an early prognostic window into infection outcome, a concept of considerable translational value for TB risk stratification.

In summary, our data demonstrate that a clinical L2-B strain of *Mtb* subverts the trained immune state established by BCG vaccination, through early suppression of antimicrobial and innate immune gene programs, disruption of key metabolic and epigenetic pathways, and selective transcription factor rewiring in lung immune cells. Crucially, we identify a context-dependent, vaccine-pressure effect evident only in BCG-vaccinated mice infected with L2-B - not in unvaccinated animals or in response to L4. Remarkably, these perturbed transcriptional signatures were not only evident in mouse lung tissue but also recapitulated in peripheral blood of humans early after exposure to L2-B *Mtb*, indicating that the mechanisms of immune evasion identified are directly relevant to the human situation. Importantly, this mechanism is not only consistent with known mycobacterial interference in chromatin and early signalling, but here we demonstrate it under BCG-driven trained immunity, within the first week of infection (or early exposure in humans), validated across species, and linked to functional impairment in the MGIA against L2-B. These findings not only offer mechanistic insight into how a hypervirulent *Mtb* strain undermines protective responses, including those mounted by trained immune cells, but also highlight the elements of BCG-induced immunity that are essential for protection against *Mtb*. The findings indicate that BCG vaccine escape is a lineage-adapted survival strategy that has major implications for TB control.

From a translational perspective, these findings argue for a re-evaluation of TB vaccine strategies. It is not sufficient to enhance immune activation alone; next-generation vaccines must also preserve the metabolic and epigenetic infrastructure that would allow durable trained immunity, and it is imperative that they are tested for efficacy against multiple different strains. Approaches such as β-glucan priming (which induces epigenetic and metabolic reprogramming in monocytes/macrophages and improves control of *Mtb* in mice)^41^, mucosal delivery of recombinant BCG strains that enhance lung-resident macrophages and engage aerobic glycolysis via mTOR or HK-1 mediated rewiring^42^, and engineered BCG overexpressing cyclic di-AMP (rBCG-DisA) which shows enhanced trained immunity signatures, epigenetic changes, and protection *in vivo*^43^ may be promising. The observation in our human household contact cohort that innate and epigenetic pathways become suppressed very early after L2-B exposure - even when individuals are still IGRA-negative - suggests that vaccine or adjunctive strategies targeting the preservation of this early trained state could act pre-emptively, possibly predicting and preventing infection before it becomes established. Such approaches might also be deployed in recent case contacts as soon as possible after exposure to boost innate immune defences and reduce the likelihood of infection.

## Materials and Methods

### Bacterial culture

Clinical isolates of *Mycobacterium tuberculosis* (Lineage 2 Beijing, Lineage 4 Euro American) were transformed by electroporation to express TdTomato fluorescence using a non-incorporating plasmid. Plasmid pTEC27 was purchased from Lalita Ramakrishnan labs^44^ (Addgene plasmid # 30182; http://n2t.net/addgene:30182; RRID:Addgene_30182). *Mtb* was grown at 37°C in Middlebrook 7H9 broth (Sigma Aldrich, St. Louis, MO, United States), supplemented with 10% Middlebrook OADC enrichment media, 0.05% Tyloxapol (Sigma Aldrich, T8761-50G). *M. bovis* BCG Pasteur TMC#1011 was cultured similarly.

### Animal studies

C57BL/6 mice were obtained from Jackson Laboratories and bred and housed at either the University of Otago or Colorado State University, in individually-ventilated cages under specific pathogen-free conditions and had ad libitum access to water and chow.

C57BL/6 mice, 6 - 8 weeks old, (n=4 males and 4 females per group for flow cytometry experiments; n=2 males per group for spatial transcriptomics) were immunised with 1×10^5^ CFU of BCG Pasteur in 40 μL of PBS intranasally while under anaesthetic restraint (intraperitoneal (i.p.) ketamine 87 mg/kg and xylazine 2.6 mg/kg). Six weeks later, mice were infected with L2-B or L4 *Mtb;* a suspension of 2 × 10^6^ CFU/ml *Mtb* in 5mL of sterile PBS was delivered by nebuliser using the Glas-Col Inhalation Exposure System (Glass-Col, Terre Haute, IN, USA). Aerosol infection deposited approximately 500-1000 CFU *Mtb* to the lungs. After infection, two mice were euthanised and whole lungs were plated onto 7H11 agar to confirm approximated CFU deposited to the lungs. Infected mice were euthanised 7-14 days after *Mtb* aerosol by i.p. injection of sodium pentobarbitone (150 mg/kg), and lungs collected for flow cytometry and spatial transcriptomic analysis.

Animal studies were carried out in accordance with the Animal Welfare Act (1999), and under the approvals of the Animal Use and Care Committee of Colorado State University, and the University of Otago Animal Ethics Committee.

### Lung harvest and single cell suspension

Lungs were perfused prior to removal by injection of PBS into the right ventricle of the heart. To obtain single-cell suspensions, lungs were cut into small pieces and incubated in Iscove’s Modified Dulbecco’s Medium (IMDM) containing 2.4 mg/mL Collagenase I (Gibco, Life Technologies) and 0.12 mg/mL DNase I (Roche Diagnostics GmbH, Mannheim, Germany) for 1 h at 37°C with 5% CO_2_. Tissue was then gently pressed through 70 μm nylon strainers. Cells were counted by haemocytometer using Trypan Blue exclusion.

### Flow cytometry

Single cell suspensions were seeded at 1.5 x 10^6^/well in 96-well V-bottom plates (Greiner Bio-One, cat#651-180). Viability staining was performed followed by Fc receptor blocking using anti-mouse CD16/32 antibody, according to **Supplementary Table 2**. Cells were then stained with a specific panel of antibodies (**Supplementary Table 2**), according to predetermined optimal dilutions in FACS buffer (PBS with 0.5% fetal bovine serum (FBS, Gibco, Life Technologies) and 2 mM EDTA). Cells were fixed in neutral-buffered Formalin (4% formaldehyde; Sigma-Aldrich). At least 300,000 events were acquired using a Cytek Aurora^™^ spectral flow cytometer (Cytek Biosciences) with ultraviolet (355nm), violet (405 nm), blue (488 nm), red (640 nm) and yellow/green (561nm) lasers. Data was analysed using manual gating set against fluorescence-minus-one controls with FlowJo 10.8.1 (Tree Star, Ashland, OR, United States) and FlowSOM unsupervised metacclustering analysis in OMIQ (Dotmatics, Boston, MA, United States), followed by identification and further immune marker analysis on clusters expressing the highest MFI for TdTomato+ *Mtb*.

### Myeloid cell enrichment and cell sorting

Lung cells were harvested and processed to single-cell suspension as described. Three lung samples were pooled to generate each final sample to increase cell numbers for cell sorting. Lymphocytes were depleted by incubating cell suspensions with APC-conjugated anti-CD3, CD5, and CD19 antibodies followed by STEMCELL Technologies’ APC Positive Selection Cocktail and EasySep Rapidspheres according to the manufacturer’s instructions. Magnet-bound cells were discarded using three rounds of magnetic separation. Depleted cell samples were then stained for flow cytometry as described.

Cells were sorted using a BD FACSAria Fusion with violet (405 nm), blue (488 nm), red (640 nm) and yellow/green (561nm) lasers. Singlet, live, GR1-, CD11c+, F4/80+ events were sorted into CD11b^hi^ and CD11b^lo^ populations and collected into IMDM supplemented with 5% FBS and 55 μM Mercaptoethanol (Gibco, Life Technologies). FMOs were used to determine gating, and 1000 events from sort products were re-acquired to assess sort purity.

### Mycobacterial growth inhibition assay

FACS-sorted CD11c+ F4/80+ CD11b^hi^ or CD11b^lo^ populations were seeded in 96-well plates and *Mtb* L2-B or L4 was introduced at a multiplicity of infection (MOI) of 0.5 (CFU:cell). Infected cultures were incubated for 3 days at 37°C with 5% CO_2_. Cells were then detached from plates and lysed by washing with sterile dH_2_0 supplemented with 0.5% Tween-80 (Sigma Aldrich) and physical scraping of the well surface. Serial 10-fold dilutions of lysates were prepared and technical replicates of six 10 μL spots per dilution were plated on 7H11 agar, incubated at 37°C. CFU were enumerated once colonies were clearly visible, 2-4 weeks after plating. The mean CFU counted from the 6 technical replicate spots per sample was calculated to generate the datapoint for each biological replicate.

### Spatial transcriptomics

Formalin-fixed, paraffin-embedded (FFPE) lung tissue blocks from mice treated with PBS, PBS + L2-B *Mtb*, PBS + L4 *Mtb*, BCG + L2-B *Mtb*, and BCG + L4 *Mtb* were soaked in ice-cold water for 2 hours. Sections (5 μm) were cut using a Leica microtome for spatial transcriptomics. Additional 10 μm sections were prepared for RNA quality assessment using the Qiagen RNeasy FFPE Kit. RNA integrity was evaluated using the DV200 metric. Sections were floated on a water bath until flat and wrinkle-free, then mounted onto the capture areas of Visium spatial gene expression slides. Slides were dried at 42°C and stored overnight in a desiccator to ensure complete dehydration.

Slides were incubated at 60°C for 2 hours, cooled, and then deparaffinised using xylene followed by graded ethanol washes (100%, 96%, 70%). Sections were stained with hematoxylin and eosin (H&E; Leica, cat# 3801570) and imaged using a KEYENCE BZ-X710 microscope. For decrosslinking, slides were treated with 0.1N HCl, followed by addition of 100 μL of TE buffer (pH 9.0), and incubated at 70°C for 1 hour in a thermocycler.

Following decrosslinking, mouse whole transcriptome probes (dual probe pairs per gene) were hybridised to tissue sections overnight at 50°C. Slides were washed, and probe ligation was performed by adding ligation enzyme and incubating at 37°C for 1 hour. Tissue permeabilisation was carried out for 40 minutes at 37°C. Subsequently, the ligated probes were extended with spatial barcodes, unique molecular identifiers (UMIs), and a partial read 1 primer.

Released cDNA probe products were collected from the slides and processed to generate sequencing-ready libraries. Sample index PCR cycle numbers were optimised using qPCR with KAPA SYBR Master Mix and primers (Roche, cat# 07959389001). Indexed libraries were amplified, purified, and size-selected using SPRIselect beads (Beckman Coulter, cat# B23318). Library quality and fragment size were assessed using the Agilent HS D1000 tape on the TapeStation system (Agilent, cat# G2991BA). Final library concentrations and molarities were quantified via qPCR, and libraries were pooled and diluted to 4 nM. Paired-end sequencing was performed on the Illumina NextSeq 500 platform at a loading concentration of 1.8 pM, using 29 cycles for read 1 and 51 cycles for read 2. Sequencing read depth was calculated based on the percentage of the capture area covered by the tissue section, multiplied by the total number of capture spots on the Visium slide (5,000 spots per capture area), and a target of 25,000 read pairs per spot.

### Computational analysis of spatial transcriptomics data

Raw BCL files generated from the Illumina sequencing run were demultiplexed using the *mkfastq* function in Space Ranger (v3.0.0, 10x Genomics). Demultiplexed FASTQ files were processed using the *count* pipeline and aligned to the mouse reference transcriptome (mm10). This pipeline performed alignment, probe collapsing, spatial barcode assignment, and gene quantification to generate feature-barcode matrices and spatial mapping information for each sample.

The filtered gene expression matrices were imported into Seurat (v4.0+) in R for downstream processing. Each sample was initially processed as an individual Seurat object. Quality control filters were applied to remove low-quality spots based on thresholds for total UMI count, number of detected genes, and mitochondrial gene content. Data normalisation and variance stabilisation were performed using Pearson residual coefficient method to reduce technical noise and correct for sequencing depth.

To enable comparative analysis across experimental groups, individual samples were merged into a single Seurat object. Prior to merging, each object was annotated with metadata including sample identity, treatment group, and capture area information. The merged sample was then normalised to harmonise expression levels across conditions.

Dimensionality reduction was performed by identifying variable features followed by Principal Component Analysis (PCA). The top PCs were used to construct a shared nearest-neighbour graph, and clusters were identified. For visualisation, Uniform Manifold Approximation and Projection (UMAP) was used to reduce high-dimensional data into two dimensions.

As Visium spots capture transcripts from multiple cells, cell type labels were interpreted as the dominant transcriptional program within each spot rather than single-cell identities. Initial annotations were assigned using the SingleR^45^ package with the MouseRNAseqData reference. To improve biological specificity in lung tissue, these labels were subsequently evaluated using a marker-based quality control approach based on canonical lineage genes. This step was used to recover immune cell signals that may be underrepresented in mixed spots due to dominant stromal or structural transcripts. Marker thresholds were defined to require consistent expression of lineage-defining genes (typically ≥2 markers per lineage) while accounting for the mixed-cell nature of spatial transcriptomic data. Marker thresholds and lineage definitions were predefined based on canonical lung cell markers and applied uniformly across all samples to minimise subjective reassignment. Cell–cell communication analysis was performed using CellChat, with interactions inferred based on refined cell type assignments.

Differentially expressed genes (DEGs) were identified using DESeq2, using group-level comparisons derived from the integrated spatial transcriptomic dataset, and the FindMarkers function was used for downstream handling and visualisation with the Seurat^46^ package. Gene set enrichment analysis was performed using clusterProfiler^47^, with both Gene Ontology^48^ (GO) and KEGG^49^ pathway analysis carried out using Mus musculus reference databases. For main figure visualisations, KEGG results were filtered at the display stage to show pathways relevant to immunological or host–pathogen processes; this filtering was applied only for figure clarity, and complete KEGG enrichment outputs for each comparison are provided in **Supplementary Tables 3 and 4**. GENIE3^50^ was used to infer group-specific regulatory networks, with candidate regulators restricted to genes detected in the dataset and classified as transcriptional regulators. This included transcription factors, chromatin modifiers, and immune-associated signalling regulators with known roles in transcriptional control. Downstream analyses focused on regulators with extensive inferred target networks, followed by GO enrichment analysis of uniquely inferred target gene sets. GO results were simplified for visualisation by grouping related terms and reducing redundancy.

### Tuberculosis household contact study

This study was embedded in a large household contact study (INFECT) which was conducted in Bandung, Indonesia, between 2014 and 2018. A detailed description of the study is found elsewhere^21^. In short, household contacts of sputum smear-positive TB patients were eligible if they were over five years of age, had no history of TB, and had lived with the adult with tuberculosis for more than five hours per week in the month prior to enrolment. They were enrolled within one week after diagnosis of the index patient and screened for TB disease using symptoms screen, chest X-ray and sputum microscopy and culture. Sociodemographic data and risk factors for *Mtb* infection were collected, including the level of exposure^21^, as measured by sleeping proximity, time spent with the index patient, and presence of cavities, and sputum mycobacterial load in the index patient.

Given the low prevalence of HIV among index patients (0.5%) and the general population at time of the study (0.2%), contacts were not tested for HIV. In Indonesia, the BCG vaccine has been included in the immunisation schedule since 1956, and vaccine coverage is high; for example, in 2013 and 2018, 87.6% and 86.9% of children under 24 months were BCG-vaccinated, respectively^51^. *Mtb* infection status of contacts was assessed by QuantiFERON-TB Gold In-Tube (QFT-GIT) IGRA, which was repeated at 14 weeks in those who were initially IGRA-negative. Based on QFT-GIT results, contacts were first classified as persistently IGRA-negative individuals and IGRA converters using the manufacturer’s cut-off value for the TBAg tube (0.35 IU/mL).

### *Mtb* strain and lineages

Sputum from the index case of the household contacts was used for *Mtb* culture and whole genome sequencing was performed on stored isolates. Lineage was determined based on a 62-single nucleotide polymorphisms barcode^52^ using TBprofiler ^53^ version 0.3.8. Only household contacts exposed to lineage 2 and lineage 4 were used in the downstream analysis. Isolates belonging to lineages 2.2.1.1, 2.2.1.2 or 2.2.2 were collectively labelled ‘Beijing’, meanwhile, lineage 2.1 were labelled as ‘non-Beijing’. The WGS data is part of a larger study, which were uploaded to the Sequence Read Archive with accession number PRJNA595834.

### Blood RNA-sequencing

Venous blood of household contacts was collected at index patient TB diagnosis, before initiation of TB treatment, into PAXgene Blood RNA Tubes (PreAnalytiX) and used for RNA-seq analysis, using the polyA tail library preparation method and single-read sequencing. Total RNA from venous blood (2.5ml) in PAXgene Blood RNA Tubes was extracted using a PAXgene blood miRNA kit (Qiagen) with the semi-automated QIAcube (Qiagen), and quantified using the LabChip GX HiSens RNA system (PerkinElmer). RNA-seq libraries were prepared using the Bioscientific NEXTflex-Rapid-Directional mRNA-seq sample preparation with the Caliper SciClone. Samples were sequenced using the NextSeq500 High Output kit V2 (Illumina) for 75 cycles.

### Statistical Analyses

Statistical analyses were performed in R version 4.4.1 (R Foundation for Statistical Computing, Vienna, Austria) using RStudio version 2024.09.0+375 (RStudio Inc., Boston, MA, United States). For spatial transcriptomics data, differentially expressed genes were identified using the FindMarkers function in Seurat. The Model-based Analysis of Single-cell Transcriptomics (MAST) hurdle model was applied for comparisons between each experimental group and PBS controls, to account for variability in gene detection and dropout typical of single-cell data. For comparisons between experimental groups the Wilcoxon Rank Sum test was used, as these groups exhibited comparable sequencing depth and gene detection profiles, making a non-parametric approach appropriate. Results were considered significant at an adjusted p-value < 0.05, unless otherwise specified.

Statistical analysis of MGIA results was performed using the Kruskal–Wallis test with Dunn’s multiple comparisons test for pairwise group comparisons. Analyses and graphs were generated using GraphPad Prism version 10.0.3 (GraphPad Software Inc., San Diego, CA, USA).

For RNA-sequencing analysis from the household contact study, only genes with transcript counts above the minimum threshold corresponding to the smallest sample size in the comparison group were included. The ‘DESeq2’^54^ package in R was used for the differential expression analysis. Due to the low number of samples (n=39), only age and IGRA status as covariates were included in the analysis and a FDR < 0.1 was considered to be statistically significant. In the analysis for specific IGRA outcome, only age was used as a covariate. Gene set enrichment analysis of the differentially expressed genes was performed using ClusterProfiler ^47^ with Gene Ontology^48^ and KEGG^49^ pathway as the reference.

## Author contributions

Conceptualisation, J.K., P.H., R.V.C., M.H.T.; Investigation, N.J.D., T.S., T.S.D.; Data analysis N.J.D., T.S., T.S.D.; Funding acquisition, J.K., R.V.C., N.J.D.; Methodology, J.K., P.H., R.V.C., M.H.T., N.J.D., T.S.; Supervision, J.K., P.H., R.V.C., M.H.T.; Writing – original draft, N.J.D., T.S.; Writing – contributions, review, editing, N.J.D., T.S., R.V.C., J.K., T.S.D., P.H., M.H.T..

## Acknowledgements

The authors kindly thank Lika Apriani (RC3ID) and Ayesha Verrall for conducting the INFECT cohort study in Bandung; Bachti Alisjahbana (RC3ID) for supporting collaborative work; Vinod Kumar (Radboudumc) for helping with transcriptional analysis of the INFECT cohort; Andres Obregon Henao for *Mtb* culture advice and CDC compliance; Megan Coleman for University of Otago Physical Containment 3 Laboratory management and MPI/CDC compliance; Amanda Hitpas and Elizabeth Creissen for technical assistance; Justin Tirados for FACS sorting of cells for MGIA; University of Colorado Anschutz for sequencing of spatial samples; Biological Research Unit at the University of Otago and Lab Animal Research at Colorado State Universir.vty for animal management.

The authors gratefully received funding from The Royal Society Te Apārangi Marsden Fund, University of Otago BMS Deans Bequest Fund, Maurice Wilkins Centre Flexible Research Seeding Programme, VALIDATE Network, Society for Mucosal Immunology, Australian and New Zealand Society for Immunology, University of Otago Division of Health Sciences.

INFECT cohort recruitment was funded by the University of Otago and Mercy Hospital (through an endowment fund and directly), Dunedin, New Zealand. Index case recruitment and investigation was part of the TANDEM project (www.tandem-fp7.eu), supported by the European Union’s Seventh Framework Program (FP7/2007–2013) under grant agreement number 305279. The IGRA (QuantiFERON) was donated by Qiagen.

---

**Supplementary Figure 1. Gene Ontology enrichment maps of host-response processes uniquely suppressed in L2-B-infected lungs relative to L4.**

Mice (n=2) were vaccinated intranasally with BCG or given PBS and challenged six weeks later with L2-B or L4 *Mtb*. Lungs were collected at day 7 post-infection, formalin-fixed, paraffin-embedded, and analysed using 10x Visium spatial transcriptomics. Left, enrichment map of GO terms uniquely suppressed in PBS + L2-B compared with PBS + L4. Right, enrichment map of GO terms uniquely suppressed in BCG + L2-B compared with BCG + L4. GO terms shown were restricted to broad, biologically relevant host-response categories, applied consistently across all groups, to highlight differences in immune, antimicrobial, metabolic, and transcriptional pathways. Nodes represent enriched GO terms, node size reflects gene set size, node colour indicates normalized enrichment score (NES), and edges connect related terms based on similarity.

**Supplementary Figure 2. Inferred regulatory strength of selected transcriptional regulators in L2-B- and L4-infected lungs.**

Mice (n=2) were vaccinated intranasally with BCG and challenged six weeks later with L2-B or L4 *Mtb*. Lungs were collected at day 7 post-infection, formalin-fixed, paraffin-embedded, and analysed using 10x Visium spatial transcriptomics. Heatmaps show inferred regulatory strength for selected transcriptional regulators across experimental groups, relative to PBS control, based on GENIE3 network analysis. Regulators shown were selected from the inferred network analysis on the basis of biological relevance.

**Supplementary Figure 3. Comparison of transcriptional programs associated with IGRA conversion versus early clearance following L2-B exposure.**

Human peripheral blood transcriptomic data from individuals who converted to IGRA positive (converters) or remained persistently uninfected (early clearers), and lung spatial transcriptomic data from BCG-vaccinated mice infected with Lineage 2 Beijing (L2-B) Mycobacterium tuberculosis were analysed. Differentially expressed genes (DEGs) were defined using p < 0.05 and |log₂ fold change| ≥ 0.5. (A) Venn diagrams showing overlap of all DEGs (left), as well as upregulated (middle) and downregulated (right) genes across human converters, early clearers, and BCG-vaccinated L2-B-infected mice. (B) Gene Ontology (GO) enrichment analysis (Biological Process) of genes upregulated in both human converters and BCG-vaccinated L2-B-infected mice, but not upregulated in early clearers. (C) GO enrichment analysis of genes uniquely upregulated in early clearers, defined as genes upregulated in early clearers but not upregulated in either converters or BCG-vaccinated L2-B-infected mice. (D) GO enrichment analysis of genes downregulated in both human converters and BCG-vaccinated L2-B-infected mice, but not downregulated in early clearers. Dot plots show the top enriched GO terms for each gene set. GO term enrichment is represented by gene ratio and adjusted p value (Benjamini–Hochberg). These analyses identify candidate transcriptional programs associated with IGRA conversion versus early clearance following L2-B exposure.

