## Supplementary Fig. 1 for "Lineage 2–Beijing *Mycobacterium tuberculosis* strains suppress BCG-trained innate immunity early after infection"

### Slide 1
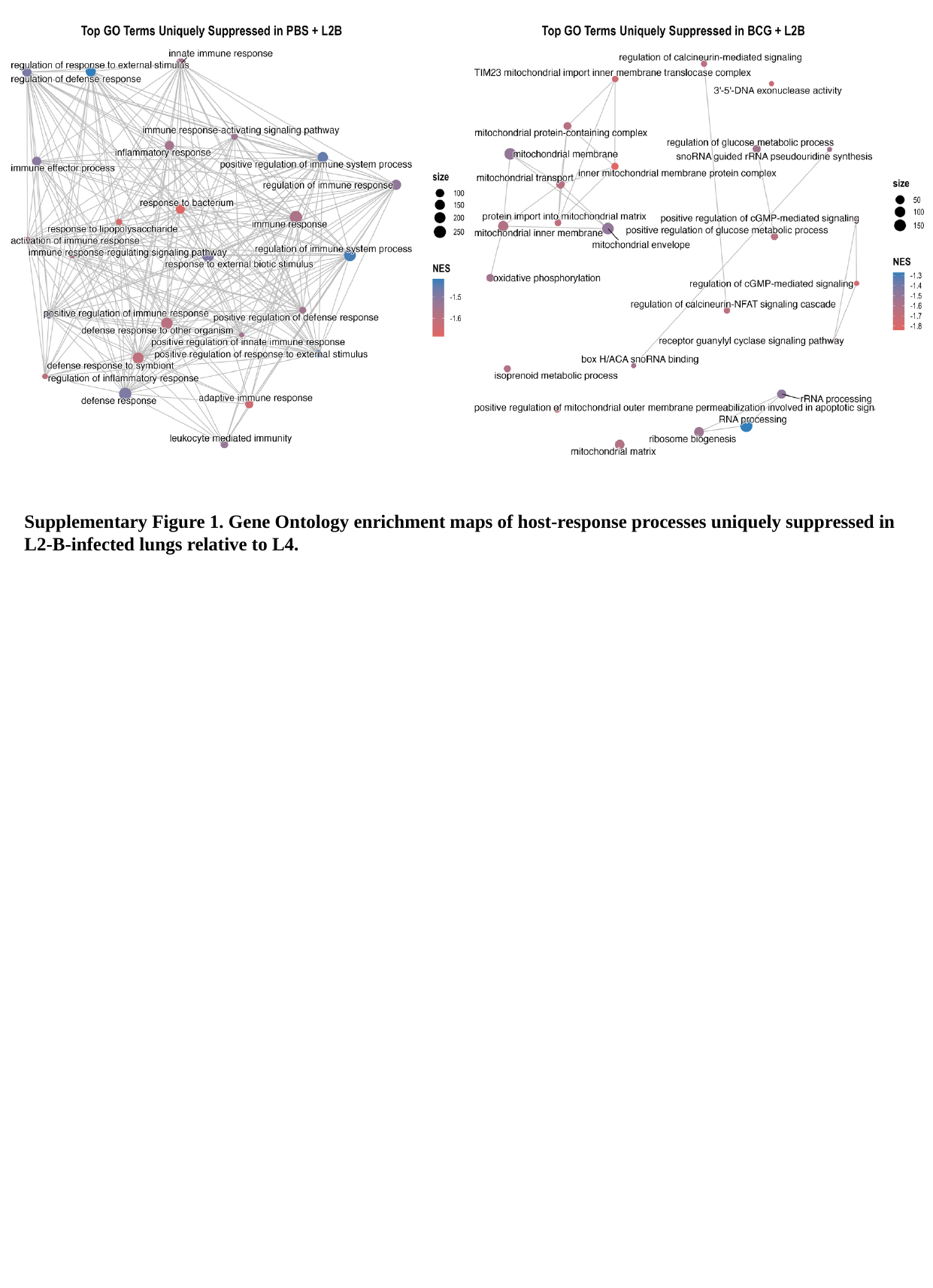

Supplementary Figure 1. Gene Ontology enrichment maps of host-response processes uniquely suppressed in L2-B-infected lungs relative to L4.
