## Supplementary Table 2 for "Lineage 2–Beijing *Mycobacterium tuberculosis* strains suppress BCG-trained innate immunity early after infection"

### Slide 1
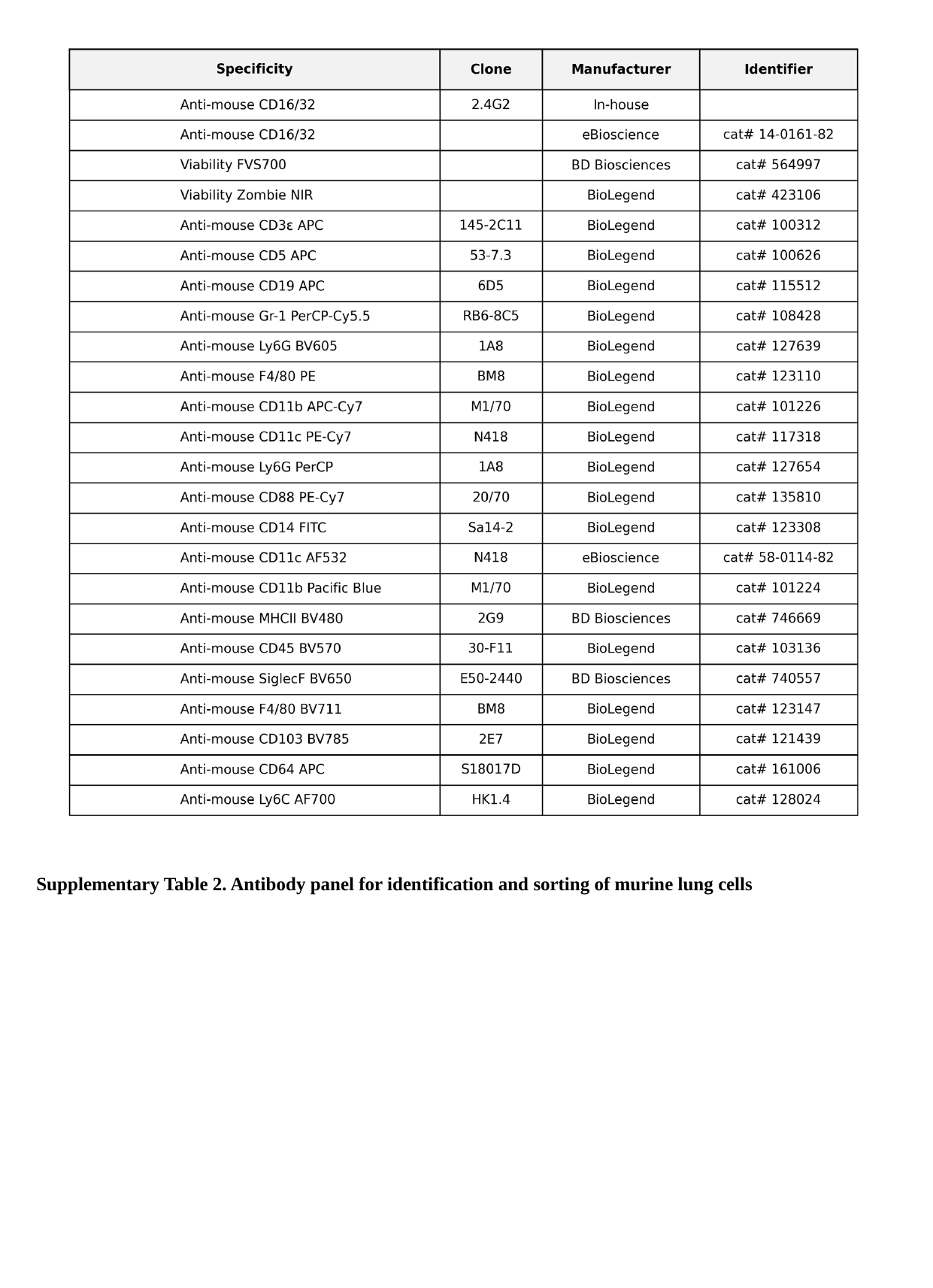

Supplementary Table 2. Antibody panel for identification and sorting of murine lung cells
