## Supplementary Fig. 3 for "Lineage 2–Beijing *Mycobacterium tuberculosis* strains suppress BCG-trained innate immunity early after infection"

### Slide 1
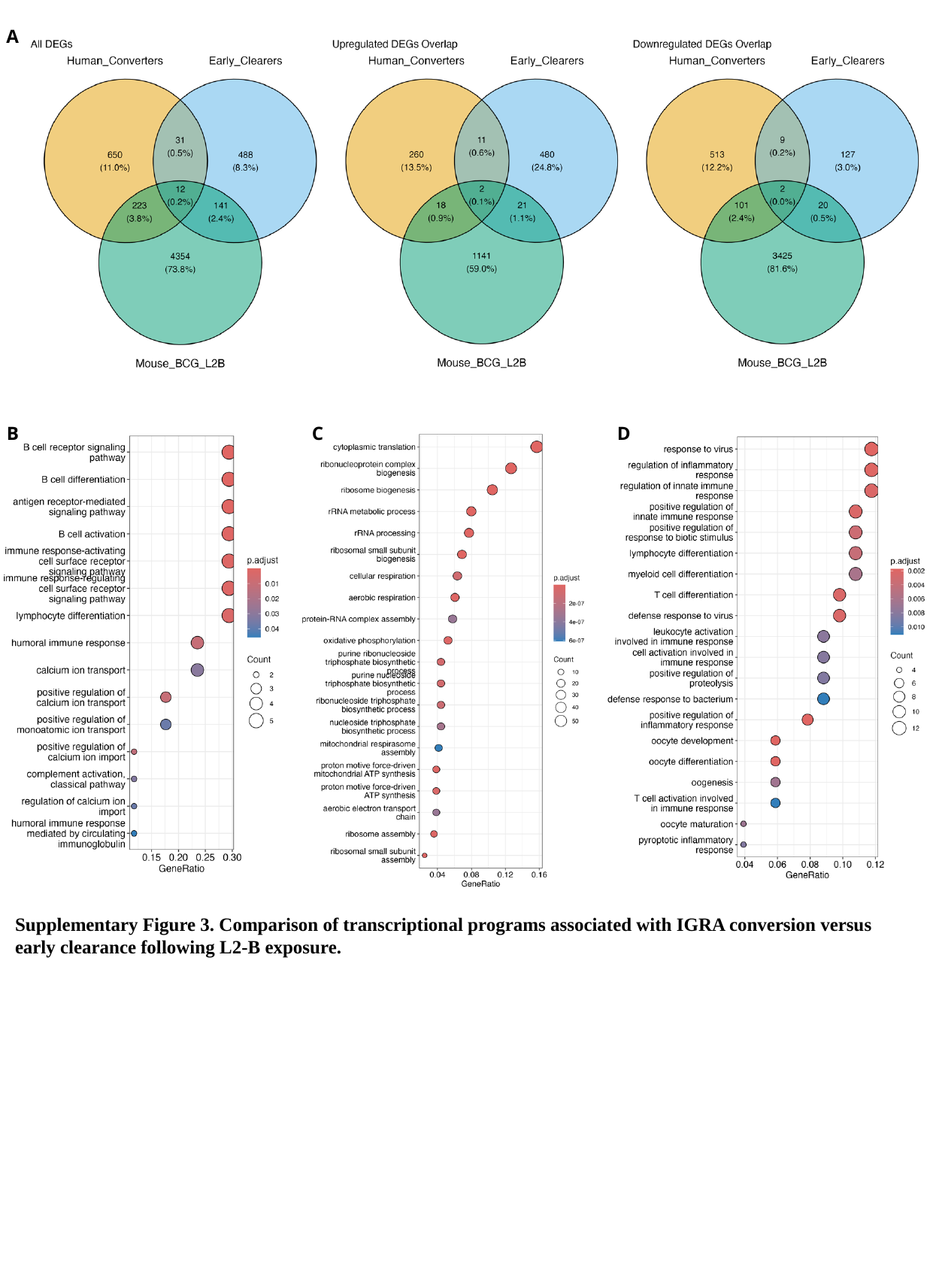

A
B
C
D
Supplementary Figure 3. Comparison of transcriptional programs associated with IGRA conversion versus early clearance following L2-B exposure.
