## Supplementary Table 1 for "Lineage 2–Beijing *Mycobacterium tuberculosis* strains suppress BCG-trained innate immunity early after infection"

### Slide 1
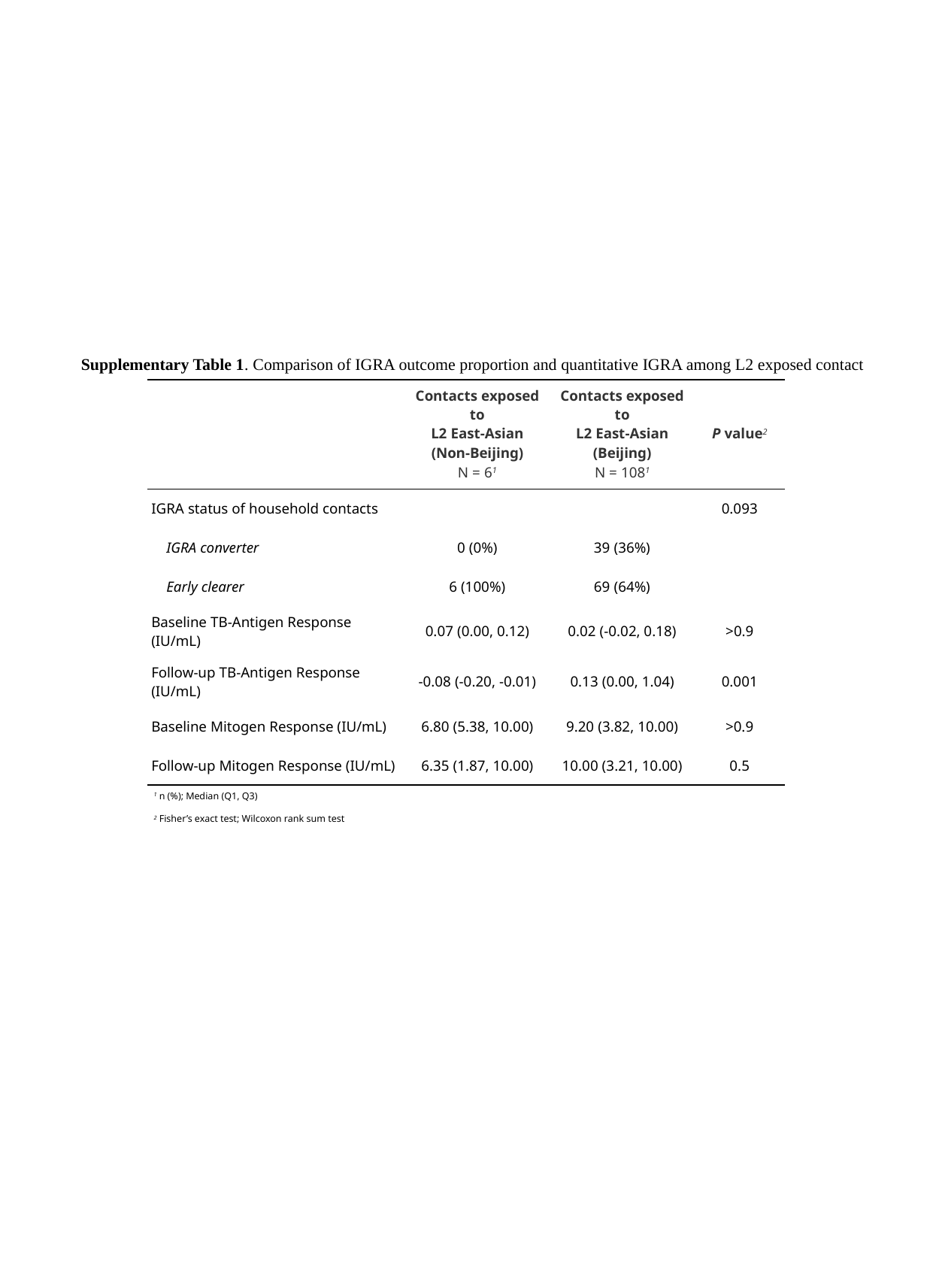

Supplementary Table 1. Comparison of IGRA outcome proportion and quantitative IGRA among L2 exposed contact
| | Contacts exposed to L2 East-Asian (Non-Beijing) N = 6^1^ | Contacts exposed to L2 East-Asian (Beijing) N = 108^1^ | P value^2^ |
| --- | --- | --- | --- |
| IGRA status of household contacts | | | 0.093 |
| IGRA converter | 0 (0%) | 39 (36%) | |
| Early clearer | 6 (100%) | 69 (64%) | |
| Baseline TB-Antigen Response (IU/mL) | 0.07 (0.00, 0.12) | 0.02 (-0.02, 0.18) | >0.9 |
| Follow-up TB-Antigen Response (IU/mL) | -0.08 (-0.20, -0.01) | 0.13 (0.00, 1.04) | 0.001 |
| Baseline Mitogen Response (IU/mL) | 6.80 (5.38, 10.00) | 9.20 (3.82, 10.00) | >0.9 |
| Follow-up Mitogen Response (IU/mL) | 6.35 (1.87, 10.00) | 10.00 (3.21, 10.00) | 0.5 |
| ^1^ n (%); Median (Q1, Q3) | | | |
| ^2^ Fisher’s exact test; Wilcoxon rank sum test | | | |
