## Supplementary Fig. 2 for "Lineage 2–Beijing *Mycobacterium tuberculosis* strains suppress BCG-trained innate immunity early after infection"

### Slide 1
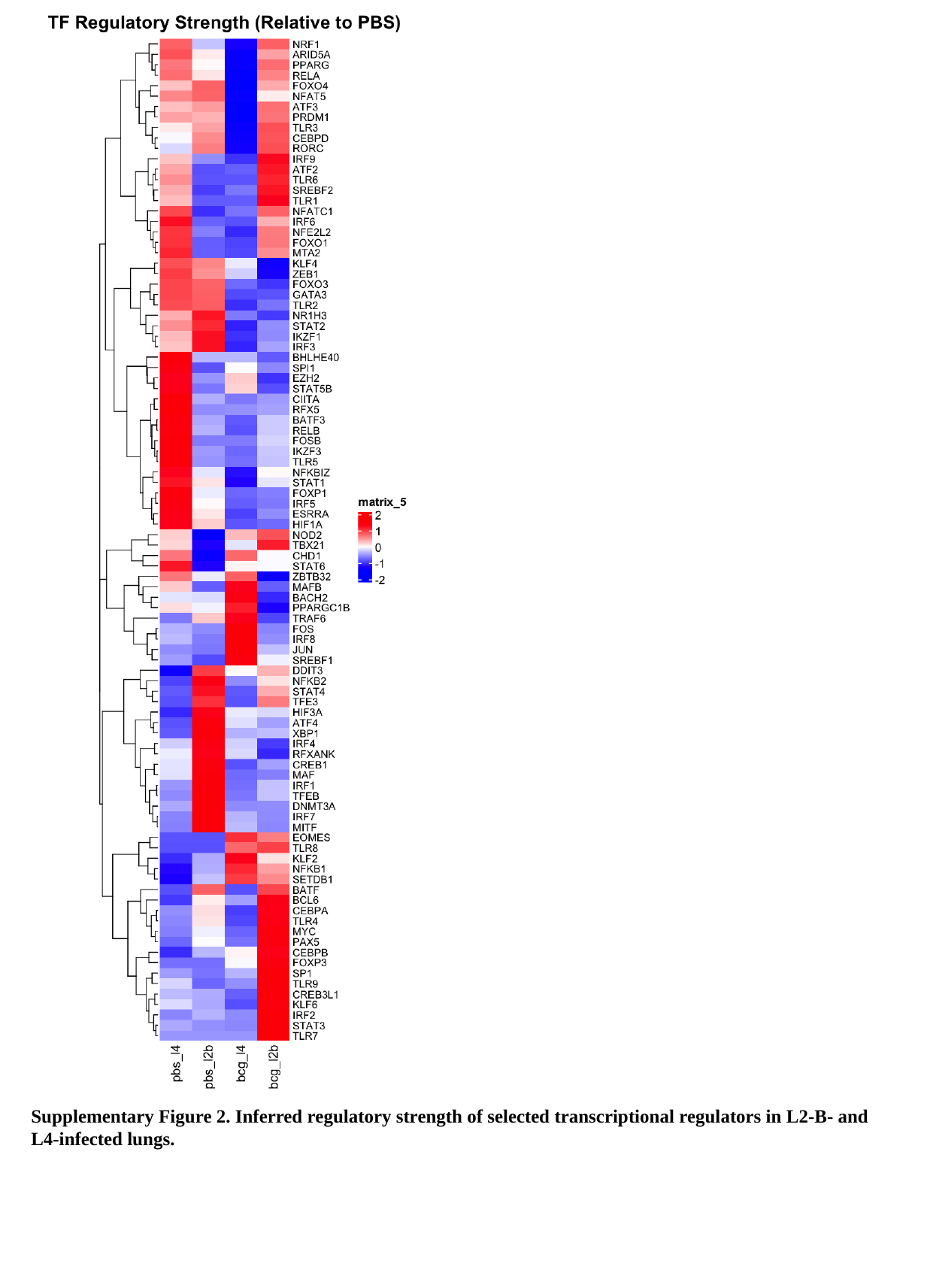

Supplementary Figure 2. Inferred regulatory strength of selected transcriptional regulators in L2-B- and L4-infected lungs.
